# MICAL-Inspired Chiral Vanadate Nanoclusters Decelerate Actin Dynamics in Live Cells

**DOI:** 10.64898/2026.06.23.733883

**Authors:** Yanan Wang, Jessica Q. Ma, Michal Sawczyk, Asli Yilmaz, E. Sumeyra Turali-Emre, Mehmet Yilmaz, Joseph Quinlan, Nicholas A. Kotov

## Abstract

Actin turnover is a fundamental cellular process essential for cell dynamics, whose control is critical for both medicine and biotechnology. However, conventional small molecules modifying actin turnover scramble the structure of actin filaments and display high cellular toxicity. MICAL enzymes oxidizing methionine (Met) residues in actin can potentially address this challenge, but their large size and multiple required cofactors make MICALs’ manufacturing and utilization difficult. Here we show that redox-active chiral decavanadate nanoclusters with tartaric acid are capable of site-selective actin modulation, mimicking MICALs, while requiring no cofactors, displaying high biocompatibility and being membrane permeable. Decavanadate nanoclusters serve as atomically precise “nano-enzymes” oxidizing three Met residues in globular actin, including Met-176; the latter inhibits the opening of the ‘backdoor’ segment and prevents depolymerization of actin filaments. The structure of actin filaments formed after nanocluster treatment revealed no structural disturbances as confirmed by cryo-electron microscopy. The biocompatibility and bioactivity of chiral decavanadate nanoclusters was demonstrated by modulation of actin in living NG108-15 cells. Taking advantage of atomically precise structure of the nanoclusters, we show that their docking into actin can be predicted computationally, indicating the possibility of programmable actin modulation using the tools of nanochemistry.

## Introduction

Actin is the key structural component of the cytoskeleton in eukaryotic cells whose regulation is essential for cellular activities.^1^ Controllable deceleration of actin dynamics is a highly sought-after capability in biomedicine and biotechnology. For instance, it is needed to reduce uncontrollable cell migration in 3D bioprinted tissues, formation of metastasis in cancer, and arrest the motility of the bacterial cells.^2^ Molecules like jasplakinolide and latrunculin stabilize and destabilize, respectively, polymerization of globular actin (i.e. G-actin) into fibrils (i.e. F-actin),^3^ but their binding leads to disorganized protein agglomerates^4^ as well as high cellular toxicity.^2,3,5^ MICAL (Molecule Interacting with CasL) enzymes offer a controllable path to deceleration of actin dynamics via selective oxidation of actin’s Met-44/47 residues, whereby promoting depolymerization of F-actin into G-actin.^6,7^ However, MICAL enzymes are difficult to manufacture and utilize due to high molecular mass, low biotechnological yield and sluggish transmembrane permeation. Furthermore, the activity of MICALs is strongly dependent on multiple co-factors and is subject to autoinhibition, which creates additional problems for their biotechnological synthesis and implementations. Thus, chemically accessible and structurally programmable compounds that can control G/F-actin turnover, while being deliverable and biocompatible, remain both fundamental and practical challenge.

Nanoparticles (NPs) that replicate enzyme activity, commonly referred to as nanozymes, have attracted significant attention due to their unique properties such as high catalytic activity, chemical robustness, and relatively low cost.^8,9^ Most reported nanozymes based on metal oxides, noble-metal composites, carbons, and metal–organic frameworks, possess characteristic sizes ranging from ∼3 to 300 nm and exhibit peroxidase-, oxidoreductase-, or catalase-like activities.^8–12^ In contrast to natural enzymes, nanozymes are generally chemo-selective rather than site- or protein-selective. This limitation arises largely from their broad size distributions, structural heterogeneity, and diverse surface chemistries. The intrinsic variability of NP structures at atomic level, together with strong nonadditive interactions^13^ not only limits selectivity but also complicates the computational prediction and modeling of NP-protein interactions. Imparting multiscale chirality and gradually controllable handedness into NPs emerged as a promising strategy to enhance the site selectivity and biological affinity of NP-protein interactions^14–17^ by mimicking lock-and-key docking of biomolecules.

Thus, we sought to create actin-targeting MICAL-like NPs to address the challenges of G/F actin regulation and complexity of MICALs’ co-factor network. Vanadium oxides were chosen as a chemical platform for these NPs because (**1**) they can reversibly access a wide range of oxidation states from +5 to +2, which is largely unavailable in other nanozymes,^18,19^ (**2**) they can be synthesized with controllable handedness using chiral surface ligands,^20^ and (**3**) vanadium compounds are biocompatible and display low systemic toxicity.^21,22^ Additionally, vanadates are structural and electronic analogues of phosphates.^23^. Since actin is also known for strong interactions with biological phosphates, such as inorganic phosphate group in different states of ionization (P_i_), ATP and ADP, ^24,25^ one may expect that vanadates of appropriate size can also dock into actin.^26^

### Synthesis and characterization of chiral decavanadate nanoclusters

We started with synthesizing V_2_O_3_ NPs stabilized by *L-* and *D*-TA (Fig. S1) in aqueous solution at pH 10.^27^ The short, oxyphilic ligands of TA promote supramolecular interactions analogous to those found in biomolecules, but absent in larger nanostructures functionalized with aliphatic thiols.^28,29^ The resulting V_2_O_3_ NPs were predominantly spherical, with an average diameter of ∼2.3 nm. Circular dichroism (CD) and ultraviolet-visible (UV-vis) spectra of V_2_O_3_ NPs recorded immediately after synthesis displayed pronounced spectral features at 400, 520, 630, and 900 nm (Fig. S1).

Importantly and unexpectedly, V_2_O_3_ NPs spontaneously convert into magic-sized nanoclusters by incubation in phosphate-buffer saline at pH 5.8, 6.4, and 7.0 at room temperature (Table S1). The chemical origin of this conversion is that NPs are destabilized at low pH by partial protonation of surface ligands.^30^ Simultaneously, acidic/neutral media favors formation of decavanadates.^31^ The conversion process (Fig. S2) was monitored by ^51^V NMR and CD spectroscopy. Because vanadium (III) from V_2_O_3_ NPs precursor is silent in ^51^V NMR, the corresponding NMR resonances appear only after three days of incubation, and growth until eight days (Fig. S3). Their gradual increase in intensity indicates that chiral nanoclusters form as a consequence of the oxidation of vanadium (III) to vanadium (V) by atmospheric oxygen and eventually formed decavanadates (Eq. S1 and S2). Decavanadates contain three crystallographically distinct vanadium atoms, which can be resolved by ^51^V NMR at - 425.7, -500.8, and -516.5 ppm from a benchmark reaction with NH_4_VO_3_ under the consistent condition as for V_2_O_3_ NPs (Fig. S4, Fig. S5). We note that the obtained dispersions exhibit ⁵¹V NMR resonance at -524.5 ppm with a shoulder near -540 ppm (Fig.S3c), which is shifted relative to the expected -516.5 ppm position observed in other decavanadates.^31^ The enhanced shielding of the vanadium nuclei causing the NMR shift is attributed to elevated electron density around the vanadium atoms arising from coordination with electron-rich, negatively charged TA ligands.

The chemical conversion of V_2_O_3_ NPs into chiral nanoclusters was also observed by CD spectroscopy because the molecular chirality of the TA ligands is transferred into chiral distortions of the decavanadate core. The typical CD peaks of V_2_O_3_ NPs gradually decreased during the incubation low pH media, while new CD features at 350 nm and 420 nm emerged (**Fig. 1a**). The attenuation of the long-wavelength CD bands (>600 nm) reflects the transition of vanadium from the +3 to the +5 state, as vanadium (V) lacks the low energy *d-d* transitions due to the transfer of *d* electrons to oxygen. The newly formed decavanadate species derived from *L-* and *D*-functionalized NPs displayed prominent absorption peaks centered at 380 nm and 480 nm and retain mirror-image CD spectra throughout the transformation, retaining their original chirality (**Fig. 1b**, **c**). The absorption bands (**Fig. 1c**) in the 380–390 nm region are assigned to core electronic transitions of the decavanadate cluster (Fig. S4f), whereas the features in the 460–480 nm range are attributed to ligand-to-metal charge-transfer transitions involving the chiral surface ligands. From a chemical standpoint, the conversion of NPs with chiral surface ligands into chiral nanoclusters with similar handedness of the core is quite remarkable considering the extensive top-to-bottom chemical changes in these nanoscale species that could have resulted in drastically diminished chirality.

**Figure 1.**
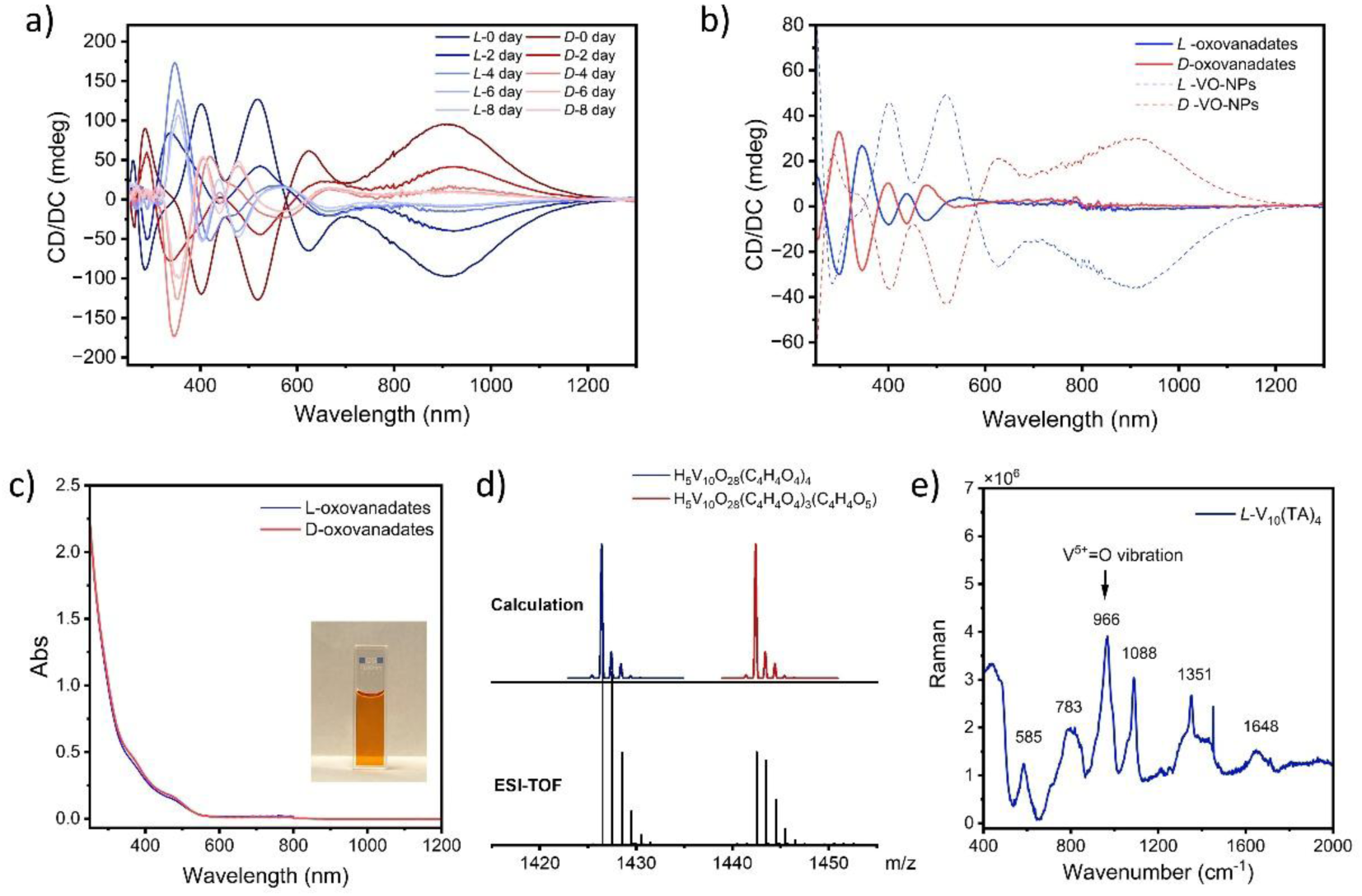
Conversion of V_2_O_3_ NPs carrying *L-* and *D*-TA into corresponding *L-* and *D*-V_10_(TA)_4_ nanoclusters. (**a**) Progression of CD spectra during incubation. (**b**) CD and (**c**) UV-Vis spectra of *L-*and *D*-V_10_(TA)_4_ and *L-* and *D*-V_2_O_3_ NPs in H_2_O. (**d**) ESI-TOF mass spectroscopy of *L*-V_10_(TA)_4_ in positive mode, flow phase H_2_O: acetonitrile 10 90, 1.0 mL/min, and calculation mass distribution for two compositions [H_5_V_10_O_28_(C_4_H_4_O_4_)_4_]^+^ and [H_5_V_10_O_28_(C_4_H_4_O_4_)_3_-(C_4_H_4_O_5_)]^+^. (**e**) Raman scattering spectra of *L*-V_10_(TA)_4_.

ESI-TOF mass spectrometry was used to characterize the product formed during the conversion of *L-*NPs, with negative and positive ionization modes examined separately. In negative mode, ESI-TOF detected only trace signals corresponding to [HV_10_O_26_]^-^ and [H_5_V_10_O_28_]^-^ species (Fig. S6). In contrast, positive mode ESI-TOF revealed intense, well-resolved signals (**Fig. 1d** and Fig. S7). The observed isotopic distributions arise from the carbon-containing organic ligand. A dominant peak at *m/z* 1426.51 corresponds to [H_5_V_10_O_28_(C_4_H_4_O_4_)_4_]^+^ (theoretical mass of 1426.41 Da) whereas higher *m/z* peaks originate from clusters containing an additional oxygen atom, assigned to [H_5_V_10_O_28_(C_4_H_4_O_4_)_3_(C_4_H_4_O_5_)]^+^ with a theoretical mass of 1442.41 Da (**Fig. 1d**). The isotopic distributions for both species are in excellent agreement with the experimental data (**Fig. 1d**). The [C₄H₄O₄] fragment (116.07 Da) arises from tartaric acid ligands bound to the V₁₀ core following the loss of hydroxyl groups at both termini. The formation of decavanadate was further confirmed by Raman spectroscopy (**Fig. 1e**). A strong band at around 966 cm^-1^, characteristic of V^5+^=O stretching vibrations, is observed and is absent in the precursor V_2_O_3_ NPs under identical pH conditions.^27,32^ Additional bands at 585 cm^-1^ and 1088 cm^-1^ were assigned to the stretching vibration of V-O bonds and typical C-O carbohydrate vibration.^27,32^ Taken together, the combined CD, ESI, and Raman spectroscopic data demonstrate that aging *L-* and *D*-V_2_O_3_ NPs under acidic conditions results in the formation of chiral decavanadate nanoclusters with four TA surface ligands. For simplicity, these nanoclusters are hereafter denoted as V₁₀(TA)₄; clusters bearing *L*- and *D*- tartaric acid ligands are referred to as *L*- and *D*-V₁₀(TA)₄, respectively.

### Chirality transfer in and conformation of V_10_(TA)_4_ clusters

To elucidate the structures of *L-* and *D*-V_10_(TA)_4_ nanoclusters and the transfer of chirality from TA to the V_10_ core (**Fig. 1a**), we performed *ab initio* density functional theory (DFT) calculations on a series of nanocluster geometries at different pH values. To probe the thermodynamic preferences underlying the experimentally observed structures and the origin of the V_10_(TA)_4_ nanoclusters (**Fig. 1d**), we first constructed a model of a TA-functionalized V_10_O_28_ core bearing four TA ligands, based on the reported crystal structure of V_10_O_28_ (**Fig. 2a**, Fig. S8).^33^ The acidity of the V_10_ nanocluster is ∼100 times higher than that of free TA (pK_a_ = 1.57 for H V O ^3−^ versus pK_a1_ = 2.98 and pK_a2_ = 4.34 for TA), leading to progressive protonation and destabilization of surface-bound ligands. This pronounced acidity promotes the gradual detachment of TA moieties from the V_10_ core as protonation increases. Accordingly, a series of partially ligand-dissociated species, namely, V_10_O_28_(TA)_2_(HTA)_2_, V_10_O_28_(TA)_1_(HTA)_3_, V_10_O_28_(HTA)_4_ and ‘naked’ V O ^6-^, were considered in our simulations (**Fig. 2a**, Fig. S8-10).

**Figure 2.**
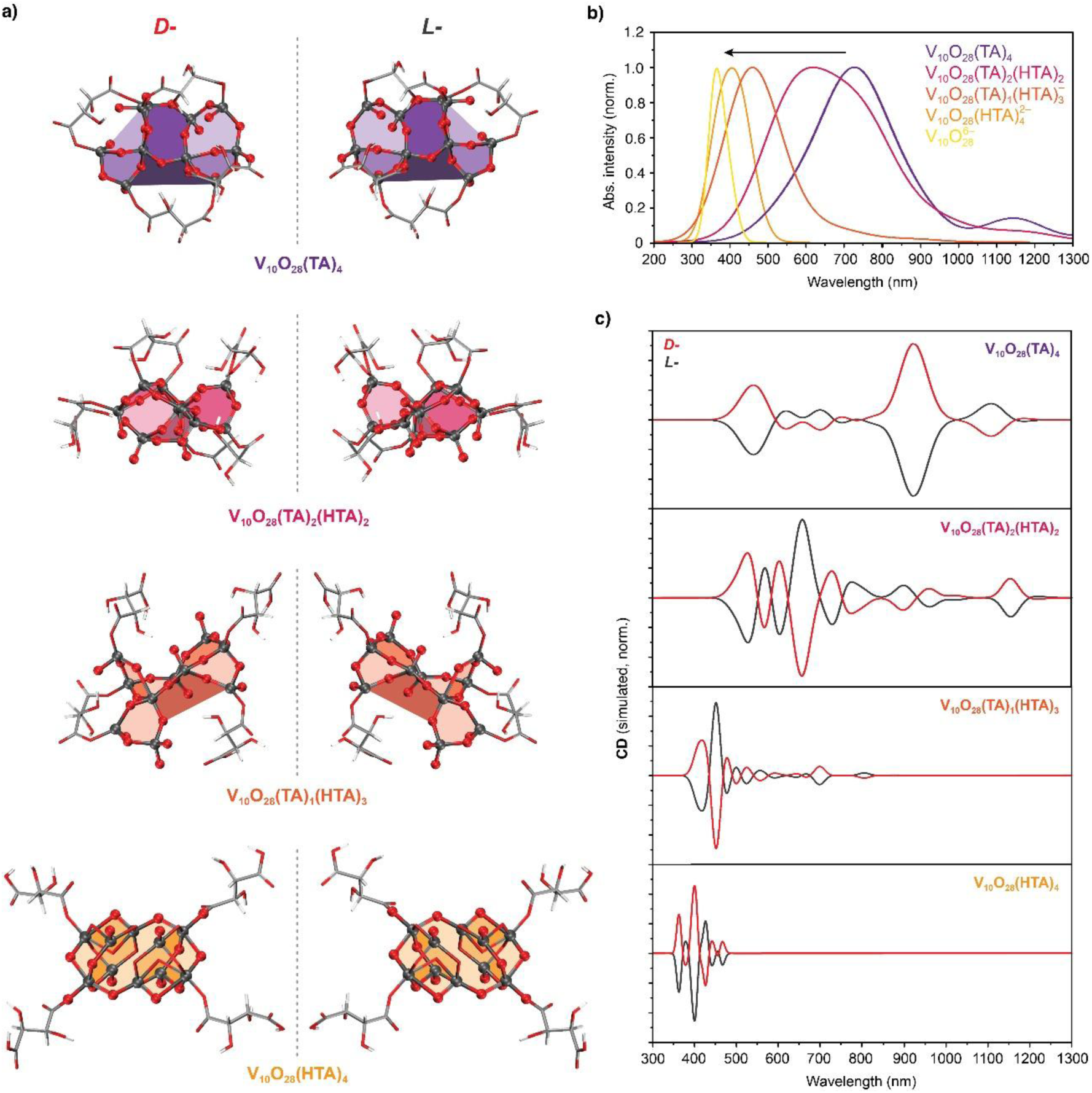
**(a)** Optimized structures of different chiral nanoclusters at different stages of protonation: V_10_O_28_(TA)_4_, V_10_O_28_(TA)_2_(HTA)_2_, V_10_O_28_(TA)_1_(HTA)_3_ and V_10_O_28_(HTA)_4_. **(b)** TD-DFT-simulated absorption spectra of chiral nanoclusters at different protonation degree of *D-*TA ligands. **(c)** Simulated CD spectra of different *D*- and *L*-V_10_ species as illustrated in **a**). HTA is a single-protonated TA residue.

To simulate the absorption and CD spectra of V O ^6-^ cores distorted by coordination with TA ligands at varying degrees of protonation, time-dependent (TD-DFT) calculations were performed (**Fig. 2b, c**). To benchmark the computational approach, we systematically evaluated the influence of different exchange-correlation functionals on the simulated CD spectra.^34^ The B3LYP functional combined with the def2-TZVP basis set and a conductor-like polarizable continuum model (CPCM) provided the most reliable and internally consistent description of the spectral features for all decavanadate species.^35^ Structural optimizations that included solvation effects revealed that progressively protonated clusters–V_10_O_28_(TA)(HTA)_3_, V_10_O_28_(HTA)_4_, and achiral V O ^6-^–were energetically more stable than the fully ligand-coordinated V_10_O_28_(TA)_4_ structure. The corresponding relative energies are *ΔE* = –17.8, –51.2, and –78.5 kJ/mol, respectively, reflecting increasing stabilization upon ligand protonation and detachment.

We calculated CD spectra of the nanoclusters across multiple protonation states (**Fig. 2c**) and observed pronounced changes in both the positions and intensities of the CD peaks. Assignment of the dominant CD peaks to HOMO-LUMO transitions indicates this such strong spectral sensitivity originates from substantial ligand-to-core (TA→core) electronic contributions, which depend critically on ligand conformation (Fig. S8-10). Notably, the sequence and relative intensities of the CD bands calculated for V_10_O_28_(TA)(HTA)_3_ closely reproduce the experimentally observed spectra of *L-* and *D*-V_10_(TA)_4_ (**Fig. 1b**, Fig. S11). As expected for TD-DFT simulations, an overall spectral shift of approximately 100 nm relative to experiment is observed.^36^

### V_10_(TA)_4_ interactions with G-actin

To investigate the interaction between G-actin and V_10_(TA)_4_, the fluorescence (FL) and UV-vis absorption spectra of native G-actin as well as the V_10_(TA)_4_-actin mixtures were recorded. The absorption bands at 280 and 200 nm (Fig. S12a) are characteristic of aromatic acid residues and peptide backbone transitions, respectively. The near-identical absorption spectra of G-actin before and after addition of V_10_(TA)_4_ indicate that the protein retains its native conformation upon interaction with the nanoclusters. After excitation at λ_ex_ = 280 nm, G-actin exhibits fluorescence emission at 340 nm (Fig. S12b) due to its tryptophan (Trp) residues (Trp-79, Trp-86, Trp-340, and Trp-356). Trp residues participate in hydrogen bonding, as well as hydrophobic and electrostatic interactions; consequently, even small conformational changes induced by interactions with V₁₀(TA)₄ nanoclusters can lead to shifts in Trp fluorescence emission.^37,38^ The emission peak of G-actin is red-shifted by interaction with any of the V_10_(TA)_4_ nanoclusters, even without changes in excitation (**Fig. 3a**, Fig. S12). At a fixed G-actin concentration of 2 μM, increasing the concentration of *L-* or *D*-V_10_(TA)_4_ up to 160 μM results in a progressive red-shift of the emission peak from ∼340 nm to ∼357 nm (**Fig. 3b**, Fig. S13). The emission maximum of pure G-actin is invariant across different excitation wavelengths (Fig. S12d) or experimental pH ranges (Fig. S12e), so the shifted emission peak of the G-actin-V_10_(TA)_4_ system must be from interactions between the actin and nanoclusters with different handedness.

**Figure 3.**
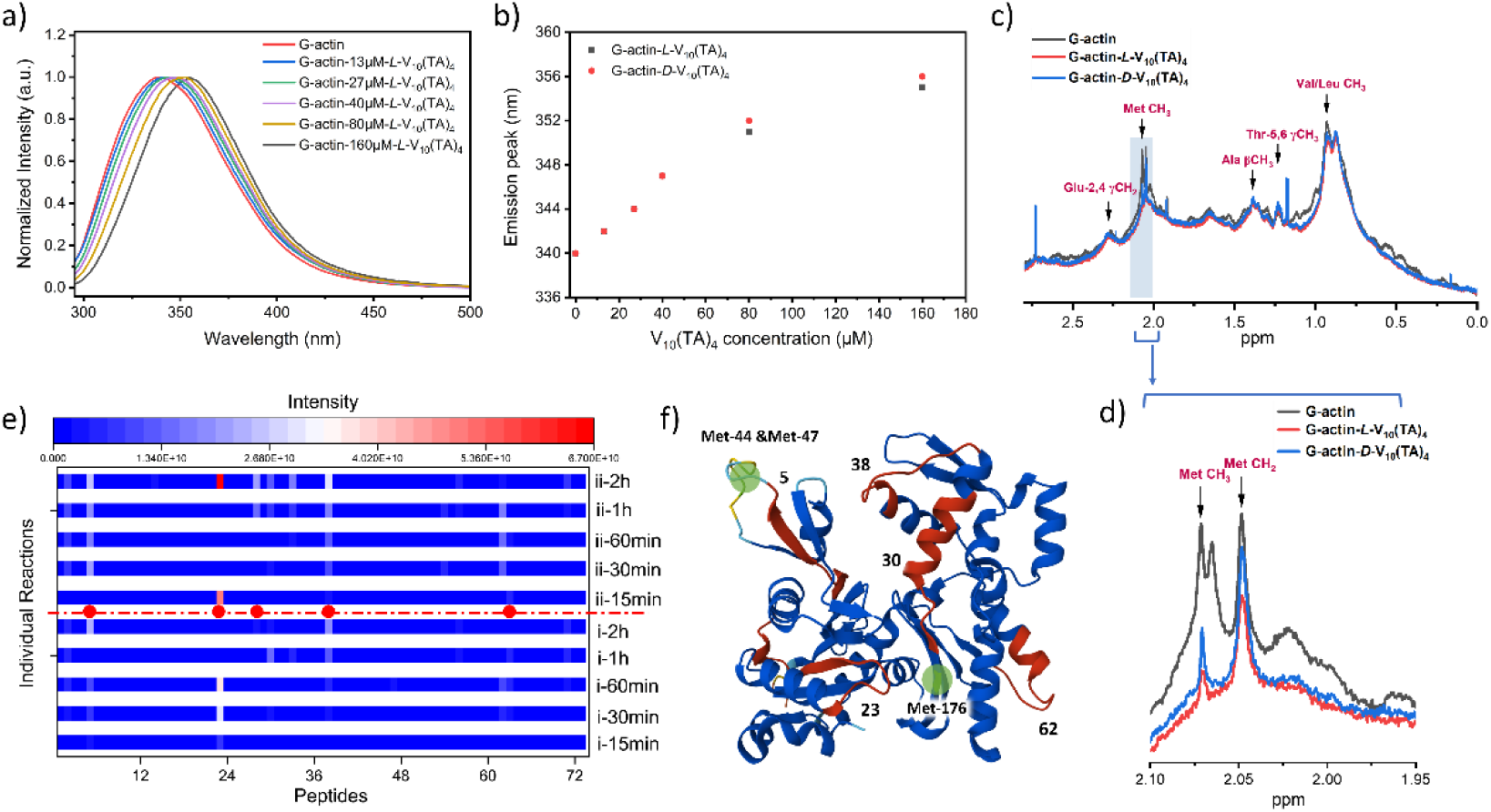
(**a**) Normalized FL emission shift of 2 μM G-actin after interacting with various concentrations of *L*-V_10_(TA)_4,_ ranging from 13 μM to 160 μM. (**b**) Spectral position of FL emission peaks of G-actin-V_10_(TA)_4_ as a function of *L-/D*-V_10_(TA)_4_ concentration. (**c**) ^1^H-NMR spectra of 30 μM G-actin (black line) plus 0.45 mM *L-* (red line) and *D-* V_10_(TA)_4_ (blue line) separately, and (**d**) zoomed-in area for the methionine (Met) region. (**e**) Representative heatmap illustrating the intensity of LC-MS/MS identified 73 trypsin-degrading peptides of G-actin (domain i), and G-actin-*L*-V_10_(TA)_4_ (domain ii) within 15 min to 2 h. G-actin and *L*-V_10_(TA)_4_ were incubated for 5 min before trypsin was added. 5 highlighted peptides are marked with red dots. A detailed peptide list is in the SI. (**f**) Peptides and residues of interest in the G-actin model. Selected peptides from LC-MS/MS were colored red, the oxidative Met residues were highlighted in green, and the G-actin structure model was extracted from AlphaFold P68135.

To experimentally probe binding sites between G-actin and V_10_(TA)_4_, a series of ^1^H NMR spectra of actin were recorded before and after incubation with *L-* and *D-*V_10_(TA)_4_ (**Fig. 3c**). The characteristic resonances from amino-acid side-chains were observed in the 0.5–2.5 ppm region (**Fig. 3c**, black trace). Additional signals arising from adenosine triphosphate (ATP) and its analogues, ADP and AMP, are also present, because of their entrapment in actin during recombinant protein production (Fig. S14). Following the addition of *L-* and *D-*V_10_(TA)_4_ to G-actin, attaining a final nanocluster concentration of 0.45 mM, resonances associated with methyl and aliphatic protons in the 1–3 ppm region decrease in intensity upon nanocluster binding in all cases (**Fig. 3c, d**). Notably, the resonances at 2.05 and 2.07 ppm, which assigned to the methylene (CH_2_) and methyl (CH_3_) groups of Met residues respectively (**Fig. 3d**), serve as sensitive spectroscopic reporters of protein-nanocluster interactions and local conformational changes.^39^ In parallel with FL measurements (**Fig. 3a**, Fig. S12), a pronounced chirality effect was observed. The complex of G-actin with *L-*V_10_(TA)_4_ exhibits the most pronounced attenuation of these Met resonances, whereas the complex of G-actin with *D*-V_10_(TA)_4_ show comparatively weaker effects (**Fig. 3d**). This differential line broadening and intensity loss indicates stronger interaction of *L-*V_10_(TA)_4_ with G-actin. Analysis of the 2.05 ppm signal yields relative shielding ratios of 18:7 for *L-* and *D-*V_10_(TA)_4_, respectively (**Fig. 3d**), which represent a significant difference for the enantiomers.

Trypsin-induced digestion of a complex of G-actin with V_10_(TA)_4_ and the protein alone was used to investigate the interaction between G-actin and nanoclusters. Analysis of the products by liquid chromatography (LC) and mass spectrometry (MS) identified a total of 73 peptides (**Fig. 3e**, Table S2). Interestingly, the formation of the G-actin-V_10_(TA)_4_ complex accelerates proteolysis of G-actin by trypsin (Fig. S15a), as evidenced by the overall increase in peptide signal intensity (**Fig. 3e**). The digestion acceleration can also be seen by taking five specific peptides in the G-actin structure (**Fig. 3f**, red).

The LC-MS/MS data indicate that three out of 14 Met residues (Met-44, Met-47 and Met-176) in G-actin are converted to methionine sulfoxide upon incubation with *L*-V₁₀(TA)₄ because of the elevated (>2) oxidation index (**Table 1**, Fig. S15b). These structural modifications are attributed to the redox enzyme-like activity of the oxovanadate nanoclusters causing Met oxidation.^19^ The resulting sulfoxide groups, together with the bound V₁₀(TA)₄ clusters, increase local polarity and promote electron withdrawal from nearby residues, causing enhanced accelerated proteolysis by trypsin (Fig. S15c). Indeed, the identified Met residues are located in the segments with enhanced MS signal intensity (**Fig. 3f**, green).

**Table 1.** Methionine (Met) side-chain oxidative modifications of actin after incubation with *L*-V_10_(TA)_4_.

| Amino acid | Oxidation index* | Mass change (Da) | Location |
| --- | --- | --- | --- |
| Met-44 | 2.4 | +15.99 | d-loop |
| Met-47 | 20.8 | +15.99 | d-loop |
| Met-176 | 2.9 | +15.99 | Pocket between SD1 and SD3 |

\*Oxidation index = Met-O @ Actin-V10 / Met-O @ pure Actin

### Selective Oxidation and Actin Turnover

MICAL enzymes typically oxidize Met-44 and Met-47 in F-actin, promoting its depolymerization into G-actin.^6,7^ Although MICALs do not oxidize Met-176, the similarity of their redox activity with V_10_(TA)_4_ is notable. The parallel becomes even more vivid when recognizing that oxidation of Met-176 controls depolymerization of F-actin into G-actin, because Met-176 is a part of the so-called ‘backdoor’ strand controlling the release of the inorganic phosphate (P_i_).^40,41^ Without the release of P_i_ formed in the process of conversion of ATP to ADP by the actin monomer units in the fibril, F-actin cannot depolymerize,^40^ which should lead to decelerated turnover and accumulation of actin fibrils.

Indeed, we observed the rapid formation of F-actin in presence of V_10_(TA)_4_ at room temperature (**Fig. 4a**). Furthermore, the accumulation of actin fibrils occurs in G-buffer that stabilizes G-actin rather than F-actin.^42^ The resulting F-actin filaments were examined by cryo-electron microscopy (cryo-EM) reconstruction (Fig. S16). The filament structure was resolved to 3.0 **Å** resolution (**Fig. 4b**, Fig. S17) with specific attention given to ADP-Ca^2+^-P_i_ state (**Fig. 4c,e,f**). A superposition of the reference model PDB:8a2y (**Fig. 4d**, green)^41^ onto our experimentally obtained filament structure (**Fig. 4d**, orange, Fig. S17c, dPDB code: 10LU, EMDB code: EMD-75276) revealed an excellent agreement. Additionally, the cryo-EM reconstruction confirmed a closed d-loop and, importantly, a closed backdoor configuration (**Fig. 4e**). The hydrogen-bonds from Arg-177 and Asn-111 residues to His-73 and Gly-74 residues from backbones create a locked polypeptide segment (**Fig. 4f**). Based on the established mechanism for P_i_ release from actin, both Arg-177 strand and Asn-111 loop need to stretch away in order to open the ‘backdoor’ and enabling P_i_ to diffuse out (**Fig. 4e**). Further inspection of the density map (**Fig. 4g**, Fig. S18) showed additional electron density on Met-176 adjacent to the sulfur atom associated with the sulfur-oxygen bond (**Fig. 4g**). The cryo-EM data confirm the mechanism of the biological activity of V_10_(TA)_4_ related to Met-176 oxidation, which stiffens the backdoor conformation, thereby slowing P_i_ release and stabilizing F-actin.

**Figure 4.**
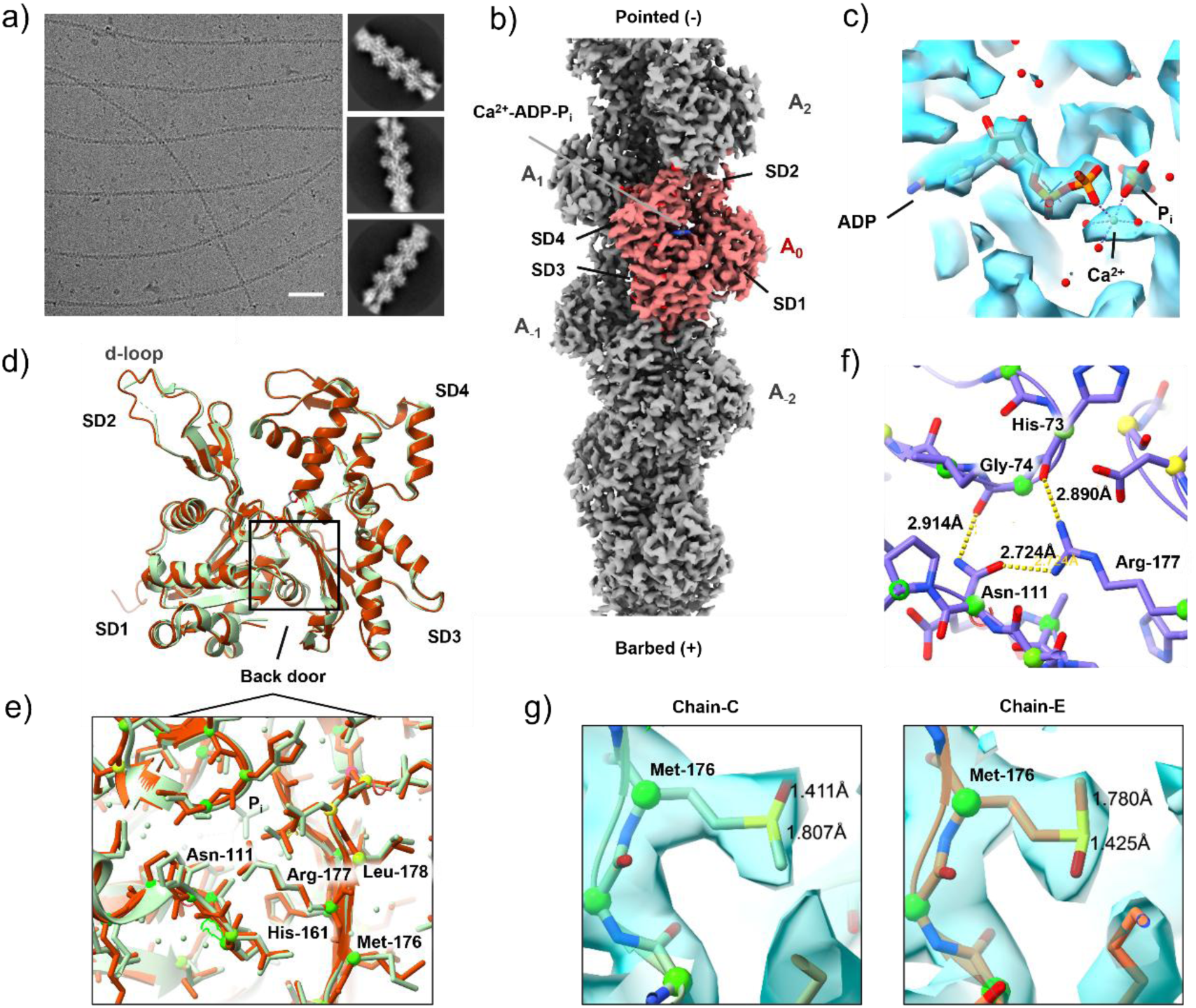
Cryo-EM reconstruction of F-actin at 3.0Å resolution. (a) Micrograph depicting actin filaments frozen in thin ice, as an example image from total database of 4,943 micrographs. Along with typical 2D classes from Cryo-EM workflow. The scale bar is 500 Å. (**b**) Bayesian polished cryo-EM density map of F-actin in the Ca^2+^-ADP-P_i_ state at 3.0 Å resolution (EMD-75276). The central subunit (A_0_, equal to Chain C in the coordinated F-actin model PDB_000010LU) is colored red, the rests (gray) are labelled based on the location along the filament. Four subdomains are marked on the central subunit (A_0_) and ADP is marked as blue inside the pocket. (**c**) Solved Cryo-EM density map of the nucleotide-binding pocket overlap with reported F-actin molecular model in Ca^2+^-ADP-P_i_ state (PDB:8a2y). (**d**) Molecular model of F-actin subunit, refined on density map of A_0_ (orange). Superimposed with model from PDB:8a2y (green). (**e**) Zoom-in on the Asn-111-Arg-177 backdoor area to feature the side chain configuration between refined molecular model (orange) and reported PDB:8a2y (green). Important residues are annotated. (**f**) Amino acid environment near residues Arg-177, Asn-111, His-73 and Gly-74 with hydrogen bonding network. **g**) Modified Met-176 residues by adding one oxygen (red) to central sulfur atom (yellow) and refined with density map inside the chain C (central subunit A_0_) and chain E (central subunit A_-2_).

One can compare the action of V_10_(TA)_4_ to conventional F-actin-stabilizing agent, such as jasplakinolide. The latter strongly perturb the native filament confirmation, creating the extensive nonspecific crosslinks and generating disorganized protein agglomerates associated with cellular toxicity.^4^ Conversely, formation of the G-actin-V_10_(TA)_4_ complex modulates actin turnover without disrupting the intrinsic structure of F-actin as indicated by the cryo-EM reconstructions in **Fig. 4**.

### Modulating Actin Turnover in Live Cells

To evaluate the bioavailability and biocompatibility of V_10_(TA)_4_, the nanoclusters were incubated with neuroblastoma×glioma hybrid NG108-15 cells.^43^ This cell line provides a particular suitable model for this study because rapid actin assembly and disassembly is essential for the neurite extension. Control of neuronal networks is also essential for 3D printed neuronal tissues^44,45^ and understanding of the metastatic behavior of brain cancer.^46^

*L*- and *D*-V_10_(TA)_4_ produced comparable effects on F-actin accumulation (**Fig. 5**) and display similar cell viability (**Fig. 5e**). Label-free refractive-index tomography was used to visualize the shape and dynamics of NG108-15 cells, including their growth-cone morphology where actin undergoes rapid turnover between fibrillar and globular states (**Fig. 5**, SI Video). Besides the changes in refractive index, the F-actin distribution in the growth cones was cross-validated by phalloidin-FITC staining (**Fig. 5a**, Fig. S19). Note that phalloidin is another actin binding agent. Unlike jasplakinolide and similarly to MICAL, phalloidin has difficulties with transmembrane transport into the cytosol and thus, used only in research. Line-profile analysis of images obtained with phalloidin-FITC tags binding to F-actin, revealed close correspondence between fluorescence and refractive index intensity for the same cellular regions (**Fig. 5b**).

**Figure 5.**
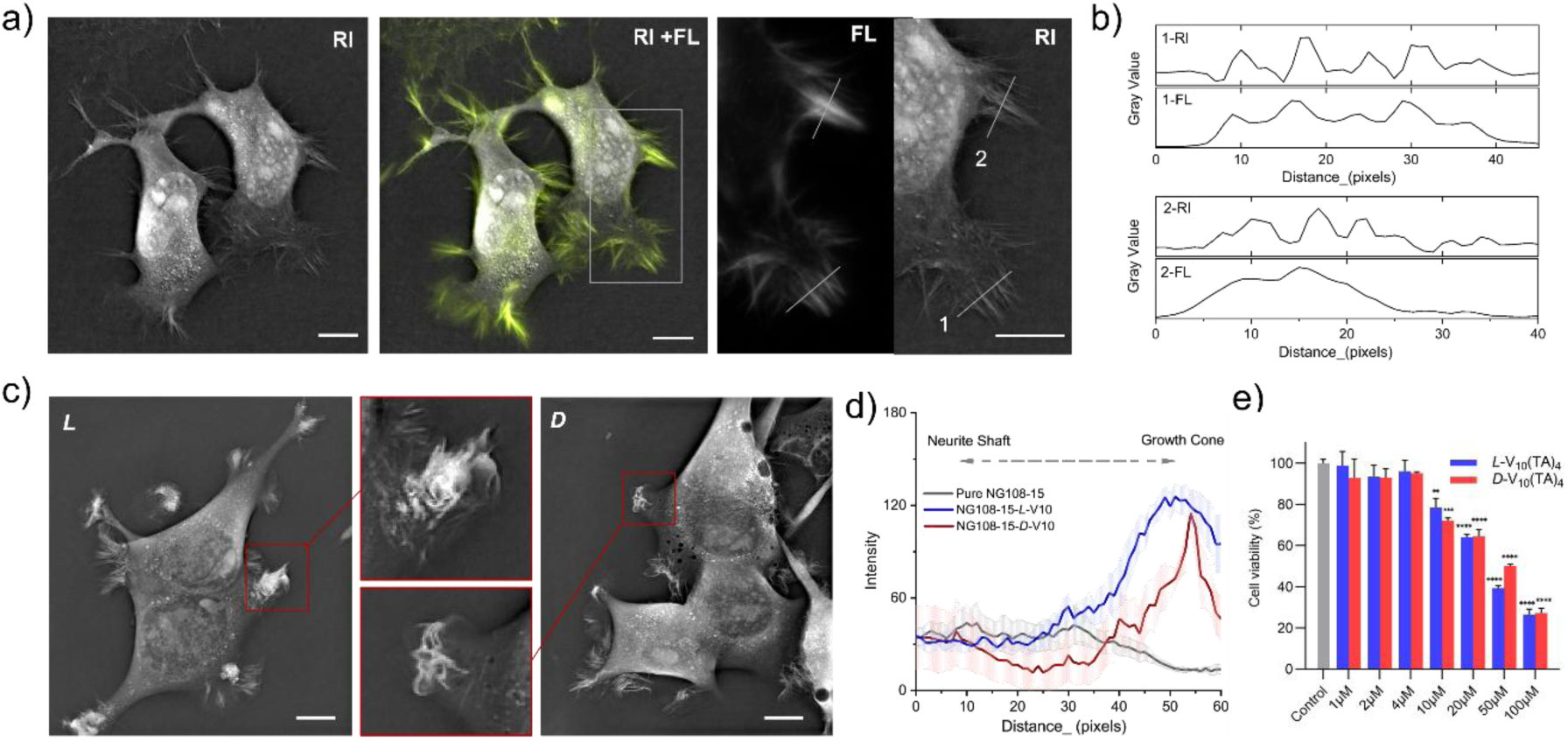
NG108-15 cell assay illustrates the F-actin accumulation at the growth cone tip after co-incubation with V_10_(TA)_4_. (a) RI and Phalloidin-FITC colocalization within NG108-15 cells. Zoomed-in views of the boxed regions were display in RI and FL channels separately with marked profile line for analysis. (b) Line-profile analysis to illustrate the correlated distribution between FL and RI signal. (c) F-actin accumulation in growth cone induced by treatment with 7uM *L*-/*D*-V_10_(TA)_4_ nanoclusters. Zoomed-in views of the red boxed regions amplified the irregular F-actin accumulation. Scale bar 10 μm. (d) Statistic intensity distribution along the 12μm line profile from neurite shaft to growth cone. Shadow area represented the statistical error from 5 individual datasets. (e) NG108-15 cell viability assay. Data are presented as mean ± SEM (n = 3 independent biological replicates). Statistical significance was defined as **p < 0.01, ***p < 0.001, ****p < 0.0001.

Without V_10_(TA)_4_, the continuous extension and retraction of filopodia was observed, which supported highly motile behavior of the neuronal cells (SI video 1-2). After incubation NG108-15 cells with *L*-/*D*-V_10_(TA)_4_ nanoclusters, a pronounced F-actin accumulation was observed at growth-cone tips, as illustrated by a higher refractive index (Fig. S20 and S21, SI video 3-5), stronger phalloidin-FITC fluorescence (Fig. S19), and increased dry-mass density. This persistent actin enrichment is consistent with F-actin stabilization and actin turnover deceleration (**Fig. 5c**) due to the inhibition of the ‘backdoor’ release of P_i_ as could be expected from **Fig. 4**. Line profiles over a 12 μm region spanning the neurite shaft to the growth cone (Fig. S22) indicate intense mass accumulation at growth cones and a significantly increased growth-cone to neurite-shaft refractive-index ratio (**Fig. 5d**) after incubation with nanoclusters.

### Computational Modeling of Nanocluster Interactions with G-actin

While the modelling of NPs interactions with biomolecules can be challenging due to their polydispersity and often uncertain surface geometry, the atomic-scale precision of V_10_(TA)_4_ enables computational modelling of their interactions with biological targets (Fig. S23).^47^ Prediction of nanocluster interactions with G-actin and potentially other proteins will facilitate their design for specific biomedical and biotechnological applications.

Because the methodologies for modelling interactions of NP and nanoclusters with proteins remain relatively underdeveloped, we compared predictions from four complementary computational frameworks: (**1**) the *ZDOCK* model, which employs rigid-body docking using a Fast Fourier Transform (FFT)-based search (Fig. S24, S25 and S26),^48^ (**2**) the *RosettaFold All-Atom* (*RFAA*)^49^ model, which combines transformer based neural networks with physics-based energy functions (Fig. S27), (**3**) the *Unified* model, which uses structural descriptors for reconfigurable nanostructures incorporating graph theoretical and chirality parameters (Fig. S28),^50^ and (**4**) the *Protein Interface Network* (*PInet*) model, which relies on surface-based representations of proteins to predict binding interfaces (Fig. S29).^51^

It is important to highlight the fundamental differences between these models. While the *Unified* model was specifically designed to transfer learning from protein-protein complexes to NP-protein complexes based on the unified structural representation, *ZDOCK*, *RFAA*, and *PInet* were not designed to accommodate structural inputs typical of NPs. Nevertheless, the atomistic precision of V₁₀(TA)₄ nanoclusters and their protein-like chemistry of particle surface coated with TA, allow *ZDOCK*, *RFAA*, and *PInet* models to be applied in a meaningful, albeit exploratory manner.

All four models also differ in their outputs. *PInet* (Fig. S29) and *Unified* (Fig. S28) estimate the relative probability of V_10_(TA)_4_ to bind to specific protein surface sites, whereas *ZDOCK* (Fig. S25) and *RFAA* (Fig. S27) evaluate the feasibility of forming discrete protein-protein complexes. In the latter case, V_10_(TA)_4_ serves as one of the proteins and G-actin is the other.

Also note that variable ligand/core chirality – a central attribute of synthetic NPs and nanoclusters --is not explicitly encoded in either *PInet* or *ZDOCK*. *RFAA* defines chirality as a binary variable (clockwise or counterclockwise), whereas *Unified* incorporates chirality through the Osipov-Pickup-Dunmur (OPD) chirality measure, which assigns continuous values reflecting both handedness and degree of chirality. Although the OPD is associated with so-called ‘chiral zeros’,^52^ incorporating chiral measures differentiating *L-* and *D-*enantiomers in one form or another remains desirable, particularly given the pronounced enantioselective differences in binding behavior observed for chiral NPs and biomolecules.^15,53–55^

Interactions between G-actin and V_10_(TA)_4_ were evaluated by all four models for variable protonation states and ligand chirality of the nanoclusters. Lower interaction probabilities and fewer binding sites were observed for complexes formed between G-actin and bare decavanadate (V₁₀O₂₈⁶⁻) versus those with V₁₀(TA)₄ nanoclusters (Fig. S24, Table S2). All the models identified similar binding sites of V_10_(TA)_4_ on G-actin but with different propensity to interact (**Fig. 6a**). Specifically, *Unified* (**Fig. 6b-c**) and *PInet* (**Fig. 6d**) predict most likely binding sites near the *N*-terminus in subdomain I and on the DNase I-binding loop (d-loop) in subdomain II. Both *RFAA* (**Fig. 6e**) and *ZDOCK* (**Fig. 6f**) predict preferential binding of V_10_(TA)_4_ within the pocket between subdomains I and III near the C-terminus of G-actin. *PInet* and *Unified* predict interactions around the d-loop, and *PInet* further identifies the ‘backdoor’ region (**Fig. 6b, 6c**). The overall predictions (**Fig. 6a**) coincide with the binding segments identified experimentally from LC-MS/MS data (**Fig. 3e-f**). In addition, prediction generated by *PInet* also recognized Met-44, Met-47, and Met-176 as residues with higher interaction probabilities with V_10_(TA)_4_ (**Fig. 6g**), which is consistent with the experiment (**Fig. 3f**). Up to 10 nanoclusters were docked into one G-actin by *ZDOCK*. The overall set of binding sites is consistent with predictions from *PInet* and *Unfield*, where the d-loop, pocked between subdomain I and subdomain III, and edge around subdomain III revealed the strongest interactions with V_10_(TA)_4_ (Fig. S26). Interestingly, the ZDOCK-scoring function for the complex between G-actin and V_10_(TA)_4_ is higher for *L-*enantiomer than that for the *D*-enantiomer of the nanocluster (Table S3), which coincides with experimental FL and ^1^H-NMR spectra of G-actin (**Fig. 3d**).

**Figure 6.**
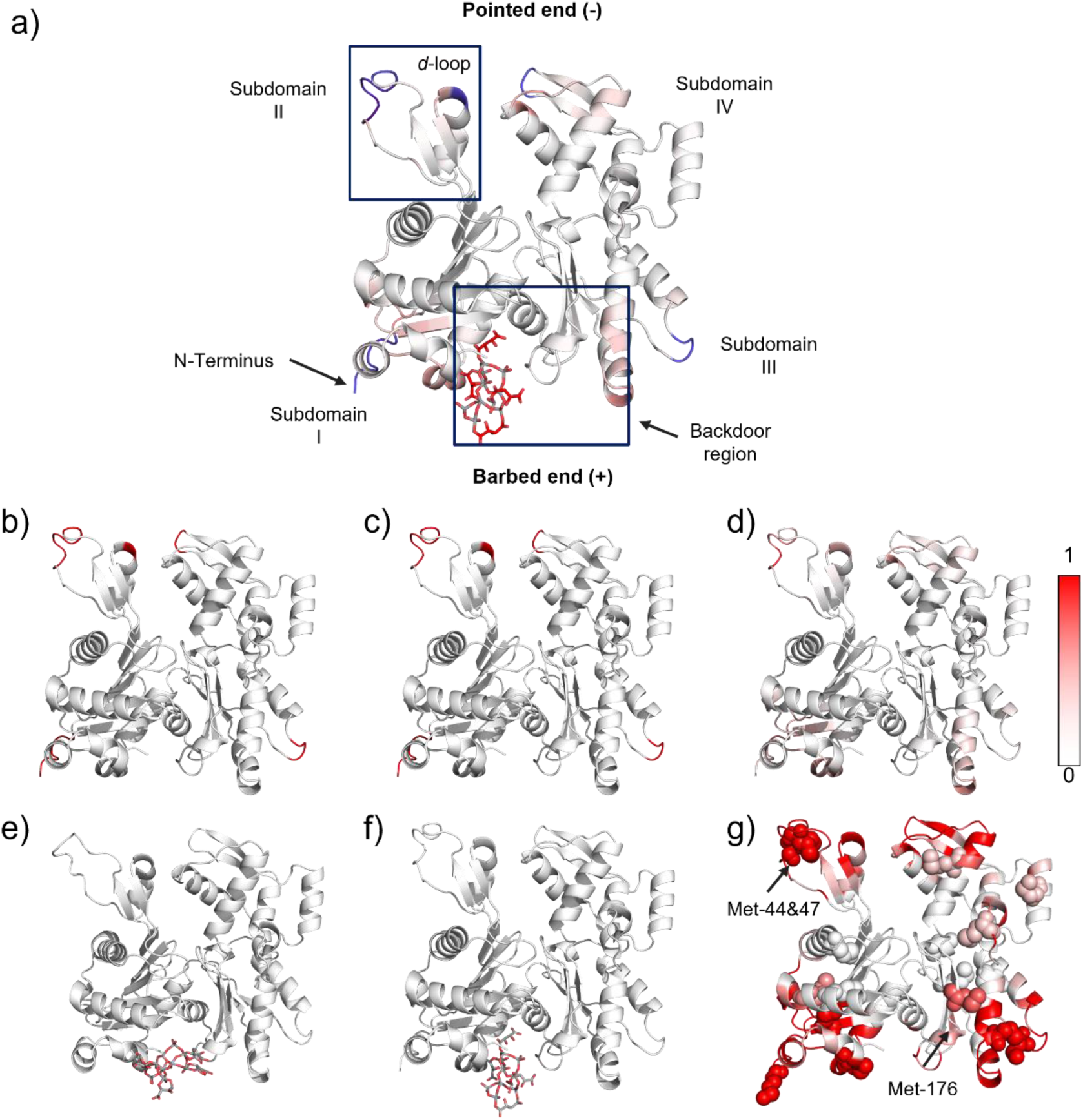
Prediction of binding sites between V_10_(TA)_4_ and G-actin. (**a**) Summarized predictive binding location. *Unified* predictions for (**b**) deprotonated *L*-V_10_O_28_(TA)(HTA)_3_, (**c**) deprotonated *D*-V_10_O_28_(TA)(HTA)_3_ with G-actin. (**d**) The interaction probability for *PInet* is normalized between 0 and 1. The color represents the predicted interaction strength between the G-actin and *L*-V_10_O_28_(TA)(HTA)_3_. Residues highlighted in red have a higher probability of interaction. (**e**) The *RFAA* predicted interaction and (**f**) the top *ZDOCK* complex between *L*-V_10_O_28_(TA)(HTA)_3_ and G-actin. (**g**) Met-focused prediction of interaction probability between *L*-V_10_O_28_(TA)(HTA)_3_ and G-actin generated using *PInet*; Met residues are shown as spheres and highlighted using a reduced maximum threshold.

## CONCLUSIONS

Inorganic analogs of MICAL enzymes were engineered based on chiral V_10_(TA)_4_ nanoclusters. They were synthesized by reorganization of chiral vanadate NPs at low pH in aqueous media, while retaining their chirality and handedness. V_10_(TA)_4_ nanoclusters selectively oxidize three Met residues in G-actin, including Met-176 controlling the “backdoor” pathway for P_i_ release. Unlike MICALs, V_10_(TA)_4_ can be easily transported through cellular membranes and do not require complex co-factors. Cryo-EM reconstruction indicate that structure of the fibrils formed in the presence of V_10_(TA)_4_ is identical to native F-actin, in contrast to the widely used actin turnover agents that result in altered structure of fibrils and protein aggregates. Cell culture testing of V_10_(TA)_4_ indicates their biocompatibility and F-actin stabilization. The latter leads to experimentally confirmed decelerated cytoskeleton remodeling, reduced dynamics of filopodia and increased F-actin accumulations in the growth cones.

To the best of our knowledge, no physiological enzyme or synthetic NPs has been reported to catalyze such biological effects, suggesting that nanoclusters can access modes of biocatalytic regulation beyond those available to natural enzymes or nanozymes. The parallels between computational predictions and experimental observations highlights the potential programmability of the V_10_(TA)_4_ nanoclusters as a framework for designing enzyme-like catalysts relevant to biotechnology and nanomedicine, that combine synthetic robustness with biomolecular selectivity.

## Data availability

All data are available in the manuscript and the supporting information. The raw data were uploaded to the server and are available for download. Code for unified structural descriptors (*Unified*) are available from the link https://github.com/jessqma/unified_fast/. The cryo-EM density map, mask map, and other EM resources and coordination files deposited at wwPDB under PDB code: PDB_000010LU, EMDB code: EMD-75276, and both entries have been approved.

## Description of Video Files

**Supplementary Video 1 | Basal growth-cone motility in live NG108-15 cells.**

Label-free time-lapse imaging of NG108-15 cells in culture medium using Nanolive 3D refractive-index tomography over 1 h, condensed to 15 s. The video captures neurite remodeling and dynamic growth-cone activity, including filopodial extension and retraction under unperturbed conditions.

**Supplementary Video 2 | The same condition as SV1, just a second example for the pure cell system.**

Basal growth-cone motility in live NG108-15 cells.

Label-free time-lapse imaging of NG108-15 cells in culture medium using Nanolive 3D refractive-index tomography <u>over 1.2h, condensed to 18 s</u>. The video captures neurite remodeling and dynamic growth-cone activity, including filopodial extension and retraction under unperturbed conditions.

**Supplementary Video 3 | *L*-V10(TA)4-induced remodeling of NG108-15 growth-cone dynamics in the first 3 hours**

Live NG108-15 cells were imaged in culture medium following incubation with final 7um *L*-V10(TA)4 nanoclusters up to 3 hours. The 3h time-lapse recording was condensed to 38s. Compared with basal growth-cone dynamics, nanocluster-treated cells showed persistent growth-cone mass accumulation and reduced dynamic remodeling of neurite-tip structures, consistent with altered actin turnover.

**Supplementary Video 4 | *D*-V10(TA)4-induced remodeling of NG108-15 growth-cone dynamics in the first 4 hours**

Live NG108-15 cells were imaged in culture medium following incubation with final 7um D-V10(TA)4 nanoclusters up to 4 hours. The 4h time-lapse recording was condensed to 60s. Compared with basal growth-cone dynamics, nanocluster-treated cells showed persistent growth-cone mass accumulation and reduced dynamic remodeling of neurite-tip structures, consistent with altered actin turnover.

**Supplementary Video 5 | *D*-V10(TA)4-induced remodeling of NG108-15 growth-cone dynamics after 4 hours**

Live NG108-15 cells were imaged in culture medium following up after the first 4 hours incubation with final 7um *D*-V10(TA)4 nanoclusters. Here another 4h time-lapse recording (condensed to 60s) illustrated the long-lasting effect from nanocluster-treated cells, with similar growth-cone mass accumulation and reduced dynamic remodeling of neurite-tip structures, consistent with altered actin turnover.

## Acknowledgements

The authors acknowledge essential support from the European Research Council via a collaborative Synergy grant: 101166855 - GAP-101166855, CHIRAL-PRO: *Handshake Complexes of Chiral Nanoparticles and Proteins*. Y.W. is thankful for the Postdoctoral Mobility project P500PN_206794 from the Swiss National Science Foundation (SNSF). This research was supported in part through computational resources and services provided by Advanced Research Computing (ARC) at the University of Michigan, Ann Arbor. The authors acknowledge the financial support of the University of Michigan College of Engineering, the Biointerfaces Institute, and the Michigan Center for Materials Characterization (MC2) for the use of the instruments and staff assistance. Cryo-EM research reported in this publication was supported by the University of Michigan Cryo-EM Facility (U-M Cryo-EM) and special thanks go to Dr. Vinson Lam and Dr. Alexandrea Shiflett, who assisted with obtaining Cryo-EM data and structure reconstruction. U-M Cryo-EM is grateful for support from the U-M Life Sciences Institute and the U-M Biosciences Initiative.

## Author contributions

N.A.K. conceptualized and guided the project. Y.W. designed and synthesized the new chiral vanadate materials and conducted experimental work and cryo-EM F-actin reconstruction J.M. carried out predictions using *Unified*, *PInet*, *ZDOCK,* and *RFAA* models. M.S. performed computational simulations and energy calculations to determine the speciation and structure of oxovanadates. A.Y and M.Y operated NG108-15 cell assay, J. Q. assisted with *PInet* predictions, E.S.T.E. assisted with the measurements on TEM and light scattering. All authors contributed to the data analysis. N.A.K., Y.W., J.M., and M.S. contributed to editing the manuscript.

## Notes

### Competing Interest Statement

The authors have declared no competing interest.

## REFERENCES

1. Park, C. E. et al. Super-Resolution Three-Dimensional Imaging of Actin Filaments in Cultured Cells and the Brain via Expansion Microscopy. ACS Nano 14, 14999–15010 (2020).

2. Bernstein, B. W., Maloney, M. T. & Bamburg, J. R. Actin and Diseases of the Nervous System. in Neurobiology of Actin (eds Gallo, G. & Lanier, L. M.) vol. 5 201–234 (Springer New York, New York, NY, 2011).

3. Pospich, S., Merino, F. & Raunser, S. Structural Effects and Functional Implications of Phalloidin and Jasplakinolide Binding to Actin Filaments. Structure 28, 437–449.e5 (2020).

4. Lázaro-Diéguez, F. et al. Dynamics of an F-actin aggresome generated by the actin-stabilizing toxin jasplakinolide. J. Cell Sci. 121, 1415–1425 (2008).

5. Spector, I., Shochet, N. R., Kashman, Y. & Groweiss, A. Latrunculins: Novel Marine Toxins That Disrupt Microfilament Organization in Cultured Cells. Science 219, 493–495 (1983).

6. Rajan, S., Terman, J. R. & Reisler, E. MICAL-mediated oxidation of actin and its effects on cytoskeletal and cellular dynamics. Front. Cell Dev. Biol. 11, 1124202 (2023).

7. Alto, L. T. & Terman, J. R. MICALs. Curr. Biol. 28, R538–R541 (2018).

8. Liang, M. & Yan, X. Nanozymes: From New Concepts, Mechanisms, and Standards to Applications. Acc. Chem. Res. 52, 2190–2200 (2019).

9. Shan, Y., Lu, W., Xi, J. & Qian, Y. Biomedical applications of iron sulfide-based nanozymes. Front. Chem. 10, 1000709 (2022).

10. Wang, D., Jana, D. & Zhao, Y. Metal–Organic Framework Derived Nanozymes in Biomedicine. Acc. Chem. Res. 53, 1389–1400 (2020).

11. Liu, Q., Zhang, A., Wang, R., Zhang, Q. & Cui, D. A Review on Metal- and Metal Oxide-Based Nanozymes: Properties, Mechanisms, and Applications. Nano-Micro Lett. 13, 154 (2021).

12. Gao, L. et al. Intrinsic peroxidase-like activity of ferromagnetic nanoparticles. Nat. Nanotechnol. 2, 577–583 (2007).

13. Silvera Batista, C. A., Larson, R. G. & Kotov, N. A. Nonadditivity of nanoparticle interactions. Science 350, 1242477 (2015).

14. Kumar, J. et al. Detection of amyloid fibrils in Parkinson’s disease using plasmonic chirality. Proc. Natl. Acad. Sci. 115, 3225–3230 (2018).

15. Chen, J. L.-Y., Pezzato, C., Scrimin, P. & Prins, L. J. Chiral Nanozymes-Gold Nanoparticle-Based Transphosphorylation Catalysts Capable of Enantiomeric Discrimination. Chem. - Eur. J. 22, 7028–7032 (2016).

16. Zhang, X., An, Z., An, J. & Tian, X. Bioinspired chiral nanozymes: Synthesis strategies, classification, biological effects and biomedical applications. Coord. Chem. Rev. 502, 215601 (2024).

17. Kotov, N. A., Liz-Marzán, L. M. & Weiss, P. S. Chiral Nanostructures: New Twists. ACS Nano 15, 12457–12460 (2021).

18. Zeng, X. et al. Vanadium Oxide Nanozymes with Multiple Enzyme-Mimic Activities for Tumor Catalytic Therapy. ACS Appl. Mater. Interfaces acsami.2c20878 (2023) doi:10.1021/acsami.2c20878.

19. Moussawi, M. A., De Azambuja, F. & Parac-Vogt, T. N. Discrete Hybrid Vanadium-oxo Cluster as a Targeted Tool for Selective Protein Oxidative Modifications and Cleavage. Angew. Chem. Int. Ed. 64, e202423078 (2025).

20. Fan, J. & Kotov, N. A. Chiral Nanoceramics. Adv Mater 32, 1906738 (2020).

21. Brenda, C.-T., Norma, R.-F., Marcela, R.-L., Nelly, L.-V. & Teresa, F. Vanadium compounds as antiparasitic agents: An approach to their mechanisms of action. J. Trace Elem. Med. Biol. 78, 127201 (2023).

22. Bakhshi Aliabad, H., et al. Vanadium complex: an appropriate candidate for killing hepatocellular carcinoma cancerous cells. Biometals 31, 981–990 (2018).

23. Rehder, D. The role of vanadium in biology. Metallomics 7, 730–742 (2015).

24. Sciortino, G., Aureliano, M. & Garribba, E. Rationalizing the Decavanadate(V) and Oxidovanadium(IV) Binding to G-Actin and the Competition with Decaniobate(V) and ATP. Inorg. Chem. 60, 334–344 (2021).

25. Schüler, H. ATPase activity and conformational changes in the regulation of actin. Biochim. Biophys. Acta BBA - Protein Struct. Mol. Enzymol. 1549, 137–147 (2001).

26. Ramos, S., Moura, J. J. G. & Aureliano, M. Actin as a potential target for decavanadate. J. Inorg. Biochem. 104, 1234–1239 (2010).

27. Shao, X. et al. Voltage-Modulated Untwist Deformations and Multispectral Optical Effects from Ion Intercalation into Chiral Ceramic Nanoparticles. Adv. Mater. 35, (2023).

28. Li, Y. & Jin, R. Seeing Ligands on Nanoclusters and in Their Assemblies by X-ray Crystallography: Atomically Precise Nanochemistry and Beyond. J. Am. Chem. Soc. 142, 13627–13644 (2020).

29. Wang, Y. & Bürgi, T. Ligand exchange reactions on thiolate-protected gold nanoclusters. Nanoscale Adv. 3, 2710–2727 (2021).

30. Kini, S., Kulkarni, S. D., Ganiga, V., T. K, N. & Chidangil, S. Dual functionalized, stable and water dispersible CdTe quantum dots: Facile, one-pot aqueous synthesis, optical tuning and energy transfer applications. Mater. Res. Bull. 110, 57–66 (2019).

31. Aureliano, M. & Crans, D. C. Decavanadate (V10O286-) and oxovanadates: Oxometalates with many biological activities. J. Inorg. Biochem. 103, 536–546 (2009).

32. O’Dwyer, C. et al. Vanadate Conformation Variations in Vanadium Pentoxide Nanostructures. J. Electrochem. Soc. 154, K29 (2007).

33. Guilherme, L. R., Massabni, A. C., Dametto, A. C., de Souza Corrêa, R. & de Araujo, A. S. Synthesis, Infrared Spectroscopy and Crystal Structure Determination of a New Decavanadate. J. Chem. Crystallogr. 40, 897–901 (2010).

34. Hirata, S. & Head-Gordon, M. Time-dependent density functional theory within the Tamm–Dancoff approximation. Chem. Phys. Lett. 314, 291–299 (1999).

35. Barone, V. & Cossi, M. Quantum Calculation of Molecular Energies and Energy Gradients in Solution by a Conductor Solvent Model. J. Phys. Chem. A 102, 1995–2001 (1998).

36. Jiang, W. et al. Emergence of complexity in hierarchically organized chiral particles. Science 368, 642–648 (2020).

37. Park, B. et al. Determination of the molecular assembly of actin and actin-binding proteins using photoluminescence. Colloids Surf. B Biointerfaces 169, 462–469 (2018).

38. Dos Santos Rodrigues, F. H., Delgado, G. G., Santana da Costa, T. & Tasic, L. Applications of fluorescence spectroscopy in protein conformational changes and intermolecular contacts. BBA Adv 3, 100091 (2023).

39. Ruschak, A. M. & Kay, L. E. Methyl groups as probes of supra-molecular structure, dynamics and function. J. Biomol. NMR 46, 75–87 (2010).

40. Oosterheert, W. et al. Molecular mechanisms of inorganic-phosphate release from the core and barbed end of actin filaments. Nat. Struct. Mol. Biol. 30, 1774–1785 (2023).

41. Oosterheert, W., Klink, B. U., Belyy, A., Pospich, S. & Raunser, S. Structural basis of actin filament assembly and aging. Nature 611, 374–379 (2022).

42. Selden, L. A., Gershman, L. C. & Estes, J. E. A kinetic comparison between Mg-actin and Ca-actin. J. Muscle Res. Cell Motil. 7, 215–224 (1986).

43. Nelson, P., Christian, C. & Nirenberg, M. Synapse formation between clonal neuroblastoma X glioma hybrid cells and striated muscle cells. Proc. Natl. Acad. Sci. 73, 123–127 (1976).

44. Joanne Wang, C., et al. A microfluidics-based turning assay reveals complex growth cone responses to integrated gradients of substrate-bound ECM molecules and diffusible guidance cues. Lab. Chip 8, 227–237 (2008).

45. Xiao, L., Mahto, S. K. & Rhee, S. W. Axon orientation by gradient of cytochalasin D inside microfluidic device. BioChip J. 6, 335–341 (2012).

46. Osswald, M. et al. Brain tumour cells interconnect to a functional and resistant network. Nature 528, 93–98 (2015).

47. Abramson, J. et al. Accurate structure prediction of biomolecular interactions with AlphaFold 3. Nature 630, 493–500 (2024).

48. Pierce, B. G., Hourai, Y. & Weng, Z. Accelerating protein docking in ZDOCK using an advanced 3D convolution library. PLOS ONE 6, e24657 (2011).

49. Krishna, R. et al. Generalized biomolecular modeling and design with RoseTTAFold All-Atom. Science 384, eadl2528 (2024).

50. Cha, M. et al. Unifying structural descriptors for biological and bioinspired nanoscale complexes. Nat. Comput. Sci. 2, 243–252 (2022).

51. Dai, B. & Bailey-Kellogg, C. Protein interaction interface region prediction by geometric deep learning. Bioinformatics 37, 2580–2588 (2021).

52. Moudgal, N., Ma, J., Turali Emre, E. S. & Kotov, N. A. Multiscale chiral zeros in biomolecules. Commun. Chem. https://doi.org/10.1038/s42004-025-01808-4 (2025) doi:10.1038/s42004-025-01808-4.

53. Xu, L. et al. Enantiomer-dependent immunological response to chiral nanoparticles. Nature 601, 366–373 (2022).

54. Sun, M. et al. Site-selective photoinduced cleavage and profiling of DNA by chiral semiconductor nanoparticles. Nat. Chem. 10, 821–830 (2018).

55. Gao, R., et al. Tapered chiral nanoparticles as broad-spectrum thermally stable antivirals for SARS-CoV-2 variants. Proc. Natl. Acad. Sci. 121, e2310469121 (2024).

