## Supplementary information for "MICAL-Inspired Chiral Vanadate Nanoclusters Decelerate Actin Dynamics in Live Cells"

### Materials and methods

***Chemicals***. Vanadium(III) chloride ($VCl_3$, 97%), *L*- and *D*-tartaric acid (TA, 99%), sodium hydroxide (NaOH, 97%), hydrochloric acid (37%), and ammonium metavanadate (99%) were purchased from Sigma-Aldrich. Actin from rabbit muscle (85% lyophilized powder, purity assessed by SDS-PAGE) was also obtained from Sigma-Aldrich and contains residual ATP, $Ca^{2+}$, and Tris buffer. Isopropanol (HPLC grade) was bought from Fisher Scientific. Water used in this study was double-distilled ($ddH_2O$). Fluorescein Isothiocyanate Labelled Phalloidin, and Triton X-100 were purchased from Sigma-Aldrich. MTT Cell Proliferation Assay Kit was bought from Cayman Chemical, USA.

***Synthesis of chiral decavanadates.*** Precursor vanadium oxide NPs were synthesized as previously reported.[1] Briefly, 0.12 g of $VCl_3$ and 0.18 g of *L*- or *D*-tartaric acid were dissolved in 30 mL of deionized water to obtain a final vanadium concentration of 0.025 M. The pH was adjusted to 10 by dropwise addition of 2.5M NaOH, and the mixture was magnetically stirred at room temperature for 2 hours. For incubation, the solution was diluted to final vanadium concentration 0.02 M using phosphate buffer (pH 5.8, 6.4, or 7.0), or dd$H_2O$. After one week of incubation, an orange solution formed, indicating the chiral nanoclusters form as a consequence of the oxidation of vanadium (III) to vanadium (V) by atmospheric oxygen and ready for other characterization. The oxidation process can be described by Equations 1 and 2:

**$V_2O_3 + O_2 <=> V_2O_5$,** (Eq. S1)

**$5\ V_2O_5 + 3\ H_2O <=> V_{10}O_{28}^{6-} + 6\ H^+$.** (Eq. S2)

***DFT and TD-DFT calculations.*** Density functional theory (DFT) and time-dependent DFT (TD-DFT) calculations were performed using the ORCA 5.0.4 software package.[2] Implicit solvation was treated using the conductor-like polarizable continuum model (CPCM) with water as the solvent.[3] Initial geometries of the VO core were adapted from the hydrated $K_6[V_{10}O_{28}]$ crystal structure reported by in Ref.[4]

TA-functionalized VO nanoclusters were constructed by covalently attaching four *D*- or *L*-TA ligands via carboxylate coordination to oxygen atoms proximal to potassium counterions in the reported crystal structure. Geometry optimizations were carried out using the B3LYP functional with the def2-TZVP basis set. TD-DFT calculations were performed on optimized structures with 50–125 excited states ($N_{roots}$), and an energy convergence tolerance of$10^{-6}$ Eh. The Tamm-Dancoff approximation was evaluated but not applied, as no statistically significant differences in results or computational efficiency were observed. To avoid unphysical charge-transfer effects between ligands and VO core, TD-DFT calculations were performed on the VO core alone, retaining the optimized geometry and a fixed total charge of –6 in all cases. Multiple combinations of functionals and basis sets were tested, including B3LYP with LANL2DZ, 6-311G(d,p) and def2-TZVP, as well as ωb97x and STEOM-DLPNO-CCSD. Although ω ιτ provided improved excited state energies and UV/Vis spectra, the B3LYP/def2-TZVP approach yielded the most consistent reproduction of experimental CD spectral features across all structures. Simulated CD peaks exhibited wavelength shifts of approximately ±30–90 nm (±0.2 to 0.7 eV), while peak polarization and overall CD patterns closely matched experimental observations.

***Simulation of CD, UV-vis, and Raman spectra.*** Simulated CD, UV–vis, and Raman spectra were generated from TD-DFT and frequency calculation outputs. Gaussian broadening was applied in Avogadro software, using a velocity representation for CD transitions.

***Chiral V10-actin interaction.*** Actin stock solutions were prepared by reconstituting rabbit muscle actin powder (1 g) in $ddH_2O$ to a concentration of 10 mg $mL^{-1}$, followed by incubation on ice for 10 min. Aliquots were snap-frozen in liquid nitrogen and stored at -80 °C. Actin samples contained no fluorescent labels or bundling proteins.

For G-actin interaction studies, *L*- and *D*-$V_{10}(TA)_4$ in pH 4 acetate buffer were added separately to G-actin and incubated for 2 minutes. The resulting mixtures were then diluted within G-buffer (5 mM Tris-HCl, pH 8.0, 0.2 mM $MgCl_2$, 0.2 mM ATP, and 0.5 mM DTT) to yield a final G-actin concentration of 2 μM, by adjusting the initial volume of nanoclusters, build complex with increasing concentrations of *L*- or *D*-$V_{10}(TA)_4$ up to 160 μM.

For actin polymerization experiments, G-actin (5 μM final concentration) was mixed with $V_{10}(TA)_4$ at a 1:20 molar ratio in modified G-buffer (5 mM Tris-HCl, 0.2 mM $Ca^{2+}$, 0.2 mM ATP, and 0.5 mM DTT, pH 6.5) and incubated at room temperature for 3 h. Samples (1mL) were ultracentrifuged at 100,000 x g for 1 h, after which 800 μL of supernatant was removed to left over the concentrated F-actin for further characterization.

***Trypsin digestion and LC–MS/MS analysis***

G-actin (5 μM) was incubated with $V_{10}(TA)_4$ (1:20 molar ratio) in modified G-buffer containing 2 mM DTT. After 5 min, trypsin was added at a 1:50 (w/w) enzyme-to-substrate ratio and samples were incubated at 37 °C. Aliquots were withdrawn at 15 min, 30 min, 1 h, and 2 h, quenched with 20% acetic acid, and placed on ice.

Samples were desalted using C18 StageTips and analyzed by LC–MS/MS using an UltiMate 3000 RSLCnano system coupled to an Orbitrap Fusion Lumos Tribrid mass spectrometer. Proteomic analysis was performed using Proteome Discoverer with the SEQUEST algorithm.

***Cryo-EM sample preparation, data collection, and processing***

Cryo-EM grids were prepared by applying 3 μL of F-actin (15–25 μM) to glow-discharged Quantifoil R2/2 Ni 300-mesh grids. Plunge freezing was performed using a Vitrobot at 20 °C and 100% humidity. Data were collected on a Thermo Fisher Krios G4i microscope (300 kV) equipped with a Gatan K3 detector and BioQuantum energy filter.

Image processing was performed using RELION 5.0. Motion correction was carried out using

MotionCor2, and CTF estimation used CTFFIND-4.1. Subsequent steps included manual particle picking, 2D and 3D classification, CTF refinement, Bayesian polishing, and post-processing, yielding a final reconstruction at 3.01 Å resolution.

An initial F-actin model (PDB:8a2y) was docked into the density map using UCSF ChimeraX and refined using ISOLDE with interactive molecular dynamics simulations.

The cryo-EM density map, mask map, and other EM resources and coordination files deposited at wwPDB under PDB code: PDB_000010LU, EMDB code: EMD-75276, and both entries have been approved.

***Spectroscopic and microscopic characterization***

CD and UV–vis spectra were recorded using a JASCO J-1700 spectrometer. ESI–TOF spectra were obtained using an Agilent 6230 system. Raman spectra were collected using a BioTools ChiralRaman-2X spectrometer (532 nm excitation).

$^{1}$H and $^{51}$V NMR spectra were recorded on a Varian VNMRS 600 spectrometer. Fluorescence measurements were performed using a Fluoromax-3 spectrofluorometer. Attempts to measure circularly polarized luminescence were unsuccessful due to negligible emission from $V_{10}(TA)_4$ at 280 nm excitation.

***NG108-15 cell assay***

NG108-15 (ECACC 88112302) cells were cultured in Dulbecco's modified Eagle's medium, DMEM, supplemented with 10% fetal bovine serum, FBS, and 1% penicillin–streptomycin at 37°C in a humidified incubator containing 5% $CO_2$. Cells were maintained below 80% confluence and passaged using 0.25% trypsin–EDTA. For imaging experiments, cells were seeded on Polymer Coverslip ibidi 35mm polymer coverslips dishes to promote cell attachment. For neurite/growth-cone experiments, cells were cultured in DMEM containing 1% FBS for at least 24 h before nanocluster treatment. Cells were then treated with different concentration of chiral decavanadate nanoclusters for further investigation. Refractive index micrographs of NG108-15 cell were recorded by Nanolive 3D Cell Explorer. For cytotoxicity evaluation, NG108-15 cells were seeded in 96-well plates at a density of approximately $5 \times 10^3$ to $1 \times 10^4$ cells per well and allowed to attach overnight. MTT assay was assessed after cell been treated with chiral decavanadate for 24h, by using an MTT Cell Proliferation Assay Kit (Cayman Chemical, USA) according to the manufacturer's instructions. Experiments were performed in three independent biological replicates, and data are presented as mean ± SEM. Statistical analyses were performed using GraphPad Prism 11.0.2 (GraphPad Software, USA). Differences among groups were evaluated by one-way ANOVA followed by Dunnett's multiple-comparisons test, with the untreated control group used as the reference. Statistical significance was defined as a two-sided $P$ value < 0.05. Significance levels are denoted as $P$ < 0.01 (**), $P$ < 0.001 (***) and $P$ < 0.0001 (****).

***Interaction prediction***

Several machine learning (ML) and docking were employed to predict interactions between $V_{10}(TA)_4$ and G-actin, leveraging similarities between protein–protein and protein–NP interfaces.[5,6] G-actin structures were obtained from the AlphaFold Protein Structure Database (AF-P68135-F1-v4),[7–10] and nanocluster structures were derived from DFT calculations.

***Protein Interface Network (PInet)***

*PInet* is based on the PointNet architecture for 3D point clouds.[10,11] Inputs include solvent-accessible surface point clouds with electrostatic potentials (from Adaptive Poisson-Boltzmann Solver, APBS) and hydrophobicity values based on the Kyte-Doolittle scale.[10] To account for orientation dependence, each protein-nanocluster pair was evaluated 1000 times using random rotations. Outputs were averaged and normalized on a 0–1 scale.

***Unified structural descriptors***

The Unified model is designed specifically for protein–nanoparticle interactions and uses combined geometric and graph-theoretical descriptors.[12] An interaction cutoff threshold of 0.001 was applied in all cases.

***ZDOCK***

Rigid-body docking was performed using ZDOCK.[13] Mulliken charges from DFT calculations were assigned to $V_{10}(TA)_4$. The top ten docking poses for both L- and D-$V_{10}(TA)_4$ consistently localized near the G-actin C-terminus between subdomains I and III. ZDOCK scores reflect shape complementarity, desolvation, and electrostatics.

***RosettaFold All-Atom (RFAA)***

RFAA inputs included the G-actin sequence (UniProt P68135) and structure data files for $V_{10}(TA)_4$.[14] While the predicted alignment error (PAE) for the actin–nanocluster interface was high (>10), the protein structure itself exhibited high confidence (PAE < 10).

**Supporting tables**

**Table S1**. Compositions of different reaction systems

| **Solvent** | | **Sodium phosphate buffer (1X)** | | | |
|---|---|---|---|---|---|
| | | 5.8 | 6.4 | 7.0 | $ddH_2O$ |
| Contents | $NH_4VO_3$ | No.1 | No.2 | No.3 | x |
| | *L*-VO-NPs | No.4 | No.5 | No.6 | No.6-1 |
| | *D*-VO-NPs | No.7 | No.8 | No.9 | No.9-1 |

**Table S2**. Identified peptides from trypsin digestion (protein-P68135) of actin

| | Sequence | Length | Mass | Start position | End position |
|---|---|---|---|---|---|
| 1 | CDEDETTALVCDNGSGLVKAGFAGDDAPR | 29 | 2925.2757 | 2 | 30 |
| 2 | AGFAGDDAPR | 10 | 975.44101 | 21 | 30 |
| 3 | AGFAGDDAPRAVFPSIVGRPR | 21 | 2155.1287 | 21 | 41 |
| 4 | AGFAGDDAPRAVFPSIVGR | 19 | 1901.9748 | 21 | 39 |
| 5 | AVFPSIVGRPR | 11 | 1197.6982 | 31 | 41 |
| 6 | AVFPSIVGR | 9 | 944.54435 | 31 | 39 |
| 7 | AVFPSIVGRPRHQGVMVGMGQKDSYVGDEAQSK | 33 | 3529.7558 | 31 | 63 |
| 8 | AVFPSIVGRPRHQGVMVGMGQK | 22 | 2350.2514 | 31 | 52 |
| 9 | AVFPSIVGRPRHQGVMVGMGQKDSYVGDEAQSKR | 34 | 3685.857 | 31 | 64 |
| 10 | PRHQGVMVGMGQK | 13 | 1423.7177 | 40 | 52 |
| 11 | HQGVMVGMGQK | 11 | 1170.5638 | 42 | 52 |
| 12 | HQGVMVGMGQKDSYVGDEAQSK | 22 | 2350.0682 | 42 | 63 |
| 13 | HQGVMVGMGQKDSYVGDEAQSKR | 23 | 2506.1693 | 42 | 64 |
| 14 | DSYVGDEAQSKR | 12 | 1353.6161 | 53 | 64 |
| 15 | DSYVGDEAQSK | 11 | 1197.515 | 53 | 63 |
| 16 | DSYVGDEAQSKRGILTLK | 18 | 1979.0324 | 53 | 70 |
| 17 | RGILTLK | 7 | 799.52797 | 64 | 70 |
| 18 | GILTLKYPIEHGIITNWDDMEK | 22 | 2585.32 | 65 | 86 |
| 19 | GILTLKYPIEHGIITNWDDMEKIWHHTFYNELR | 33 | 4082.0513 | 65 | 97 |
| 20 | YPIEHGIITNWDDMEKIWHHTFYNELR | 27 | 3456.635 | 71 | 97 |
| 21 | YPIEHGIITNWDDMEK | 16 | 1959.9037 | 71 | 86 |
| 22 | IWHHTFYNELR | 11 | 1514.7419 | 87 | 97 |
| 23 | VAPEEHPTLLTEAPLNPK | 18 | 1955.0364 | 98 | 115 |
| 24 | VAPEEHPTLLTEAPLNPKANREK | 23 | 2553.3551 | 98 | 120 |
| 25 | VAPEEHPTLLTEAPLNPKANR | 21 | 2296.2175 | 98 | 118 |
| 26 | EKMTQIMFETFNVPAMYVAIQAVLSLYASGR | 31 | 3507.7604 | 119 | 149 |
| 27 | MTQIMFETFNVPAMYVAIQAVLSLYASGR | 29 | 3250.6229 | 121 | 149 |
| 28 | TTGIVLDSGDGVTHNVPIYEGYALPHAIMR | 30 | 3195.6023 | 150 | 179 |
| 29 | TTGIVLDSGDGVTHNVPIYEGYALPHAIMRLDLAGR | 36 | 3820.957 | 150 | 185 |
| 30 | LDLAGRDLTDYLMK | 14 | 1622.8338 | 180 | 193 |
| 31 | LDLAGRDLTDYLMKILTER | 19 | 2235.1933 | 180 | 198 |
| 32 | LDLAGRDLTDYLMKILTERGYSFVTTAER | 29 | 3346.7231 | 180 | 208 |
| 33 | DLTDYLMK | 8 | 997.47903 | 186 | 193 |
| 34 | DLTDYLMKILTER | 13 | 1609.8385 | 186 | 198 |
| 35 | DLTDYLMKILTERGYSFVTTAER | 23 | 2721.3684 | 186 | 208 |

| | | | | | |
|---|---|---|---|---|---|
| 36 | ILTERGYSFVTTAER | 15 | 1741.8999 | 194 | 208 |
| 37 | ILTERGYSFVTTAEREIVR | 19 | 2239.1961 | 194 | 212 |
| 38 | GYSFVTTAER | 10 | 1129.5404 | 199 | 208 |
| 39 | GYSFVTTAEREIVR | 14 | 1626.8366 | 199 | 212 |
| Continued Table 3 | | | | | |
| 40 | GYSFVTTAEREIVRDIKEK | 19 | 2240.1801 | 199 | 217 |
| 41 | GYSFVTTAEREIVRDIK | 17 | 1983.0425 | 199 | 215 |
| 42 | EIVRDIKEK | 9 | 1128.6503 | 209 | 217 |
| 43 | EIVRDIK | 7 | 871.51272 | 209 | 215 |
| 44 | DIKEKLCYVALDFENEMATAASSSSLEK | 28 | 3091.473 | 213 | 240 |
| 45 | EKLCYVALDFENEMATAASSSSLEK | 25 | 2735.267 | 216 | 240 |
| 46 | LCYVALDFENEMATAASSSSLEK | 23 | 2478.1294 | 218 | 240 |
| 47 | SYELPDGQVITIGNERFR | 18 | 2093.0542 | 241 | 258 |
| 48 | SYELPDGQVITIGNER | 16 | 1789.8846 | 241 | 256 |
| 49 | FRCPETLFQPSFIGMESAGIHETTYNSIMK | 30 | 3433.6145 | 257 | 286 |
| 50 | CPETLFQPSFIGMESAGIHETTYNSIMK | 28 | 3130.445 | 259 | 286 |
| 51 | CDIDIRK | 7 | 861.43784 | 287 | 293 |
| 52 | CDIDIRKDLYANNVMSGGTTMYPGIADR | 28 | 3088.4416 | 287 | 314 |
| 53 | CDIDIRKDLYANNVMSGGTTMYPGIADRMQK | 31 | 3475.6357 | 287 | 317 |
| 54 | KDLYANNVMSGGTTMYPGIADR | 22 | 2373.1093 | 293 | 314 |
| 55 | KDLYANNVMSGGTTMYPGIADRMQK | 25 | 2760.3033 | 293 | 317 |
| 56 | DLYANNVMSGGTTMYPGIADR | 21 | 2245.0144 | 294 | 314 |
| 57 | DLYANNVMSGGTTMYPGIADRMQK | 24 | 2632.2084 | 294 | 317 |
| 58 | DLYANNVMSGGTTMYPGIADRMQKEITALAPSTMK | 35 | 3774.8089 | 294 | 328 |
| 59 | MQKEITALAPSTMK | 14 | 1547.8051 | 315 | 328 |
| 60 | MQKEITALAPSTMKIK | 16 | 1788.9842 | 315 | 330 |
| 61 | MQKEITALAPSTMKIKIIAPPER | 23 | 2565.4386 | 315 | 337 |
| 62 | EITALAPSTMK | 11 | 1160.6111 | 318 | 328 |
| 63 | EITALAPSTMKIK | 13 | 1401.7901 | 318 | 330 |
| 64 | EITALAPSTMKIKIIAPPER | 20 | 2178.2446 | 318 | 337 |
| 65 | EITALAPSTMKIKIIAPPERK | 21 | 2306.3396 | 318 | 338 |
| 66 | IKIIAPPER | 9 | 1035.6441 | 329 | 337 |
| 67 | IKIIAPPERK | 10 | 1163.739 | 329 | 338 |
| 68 | IIAPPER | 7 | 794.46504 | 331 | 337 |
| 69 | IIAPPERK | 8 | 922.56 | 331 | 338 |
| 70 | YSVWIGGSILASLSTFQQMWITK | 23 | 2615.3458 | 339 | 361 |
| 71 | QEYDEAGPSIVHR | 13 | 1499.7005 | 362 | 374 |
| 72 | QEYDEAGPSIVHRKCF | 16 | 1877.873 | 362 | 377 |
| 73 | QEYDEAGPSIVHRK | 14 | 1627.7954 | 362 | 375 |

Note: Here the G-actin structure sequence applied from AlphaFold P68135-F1-V6, which the sequence length is 377, for residue number labelled within the structure according to after remove the first two amino acid to normalized the residue number to general literatures recording with sequence length 375.

**Table S3**. ZDOCK scores (ZDOCK) and PAE (RFAA) for enantiomers of $V_{10}(TA)_4$

| | **ZDOCK Score** | **Interaction PAE** | **Protein PAE** |
|---|---|---|---|
| *D*-$V_{10}O_{28}(TA)_1(HTA)_3$ deprotonated | 1125.912 | 27.586 | 4.882 |
| *D*-$V_{10}O_{28}(TA)_1(HTA)_3$ protonated | 1088.565 | 27.661 | 4.765 |
| *L*-$V_{10}O_{28}(TA)_1(HTA)_3$ deprotonated | 1212.035 | 27.490 | 4.784 |
| *L*-$V_{10}O_{28}(TA)_1(HTA)_3$ protonated | 1097.64 | 26.456 | 4.876 |
| Pure $V_{10}O_{28}$ | 949.285 | 26.850 | 4.949 |

Note: The ZDOCK score considers pairwise statistical potential, shape complementarity, desolvation, and electrostatics to evaluate the relative likelihood of docking. For any given configuration, L-$V_{10}(TA)_4$ has a higher ZDOCK score compared to D-$V_{10}(TA)_4$.

High interaction predicted aligned error (PAE) suggests that RFAA is not a suitable model for predicting interactions between actin and $V_{10}(TA)_4$ nanoclusters. This is also observed in the overlap between the protein and nanocluster structures in the docked complex, which is physically impossible **(Fig S27)**.

**Supporting figures**

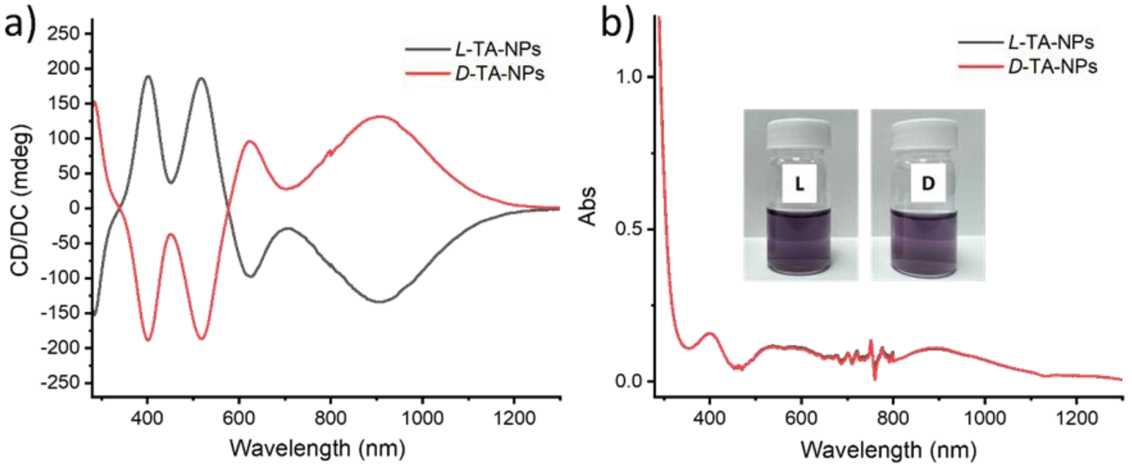


**Fig. S1 | Spectroscopic characterization of chiral $V_2O_3$ nanoparticles synthesized at pH 10**. (**a**) Circular dichroism (CD) spectra and (**b**) UV–vis absorption spectra of *L*- and *D*-$V_2O_3$ nanoparticles recorded in $H_2O$ at a final vanadium concentration of 0.02 M; insets show photographs of *L*- and *D*-$V_2O_3$ nanoparticle dispersions under visible light.

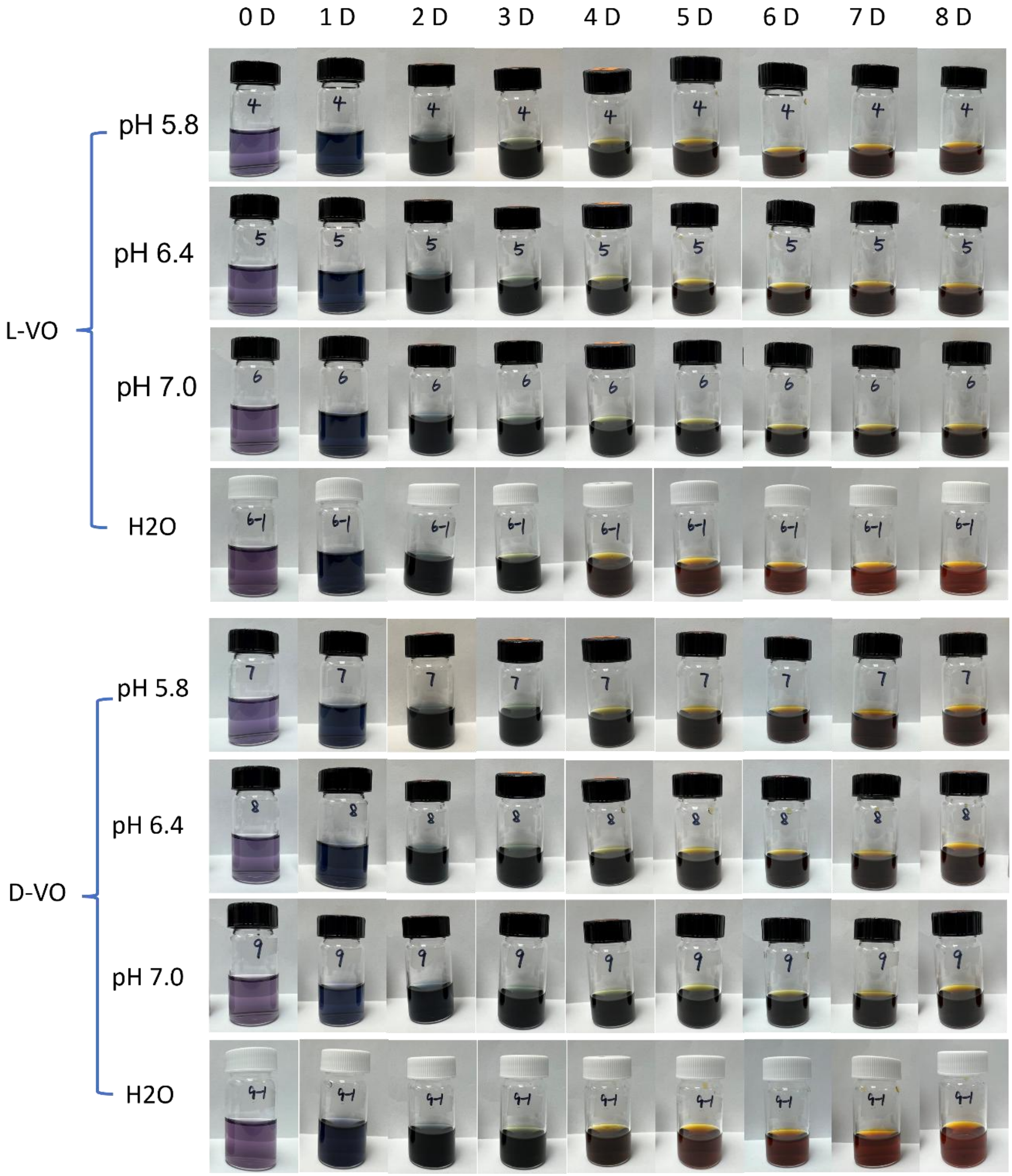


**Fig. S2 | Time-dependent evolution of chiral $V_2O_3$ nanoparticles under different pH conditions**. Chiral *L*- and *D*-$V_2O_3$ nanoparticle dispersions incubated in buffers of different pH and in pure $H_2O$ over time periods of up to 8 days.

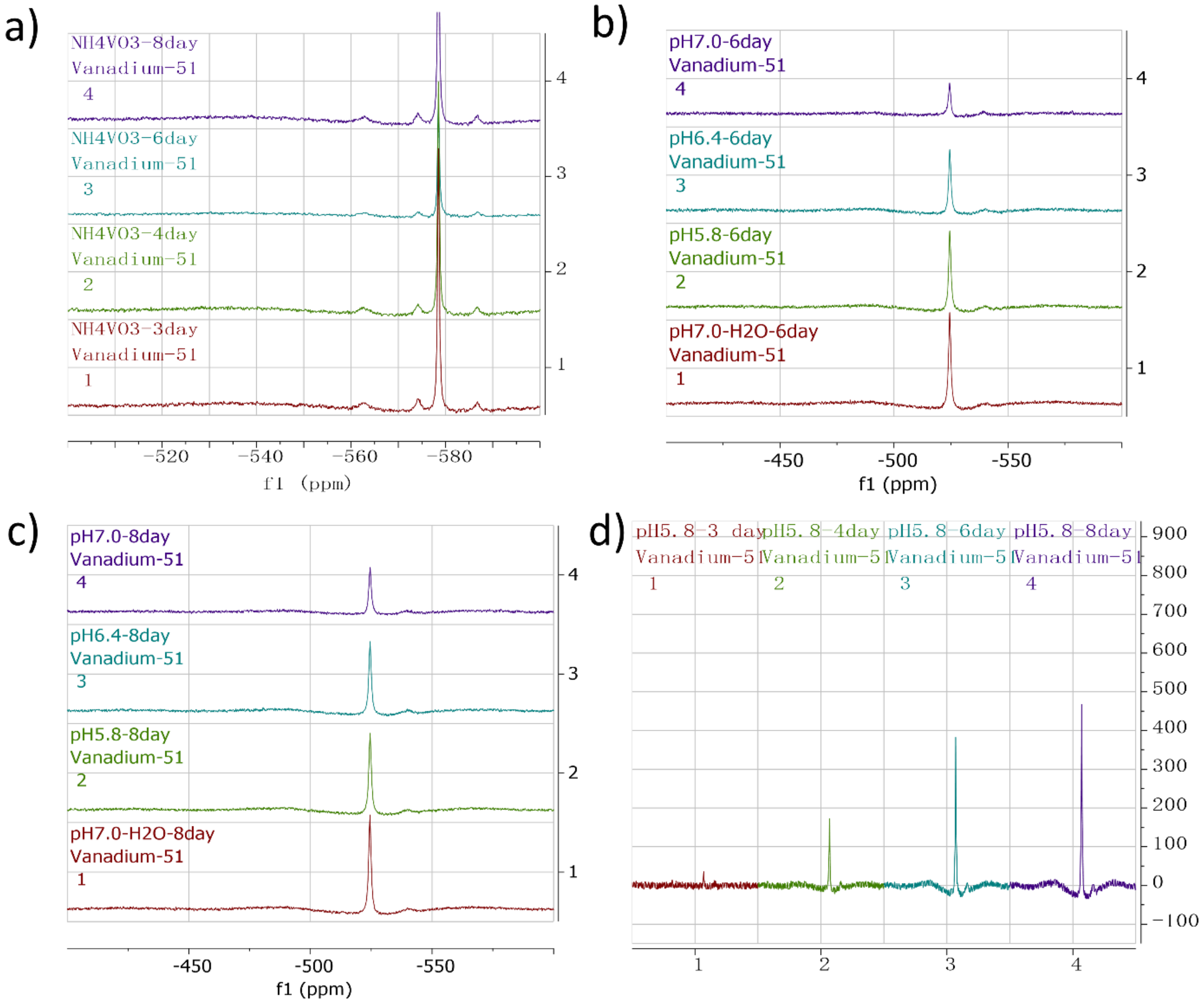


**Fig. S3 | $^{51}$V NMR analysis of vanadate species as a function of pH and incubation time.** (**a**) Dispersions of vanadate nanoparticles prepared from $NH_4VO_3$ after 3–8 days of incubation. (**b**, **c**) Dispersions of *L*-$V_2O_3$ nanoparticles incubated at different pH values after 6 days (b) and 8 days (c). (**d**) Time-dependent evolution of $^{51}$V NMR spectra for dispersions of *L*-$V_2O_3$ nanoparticles incubated at pH 5.8.

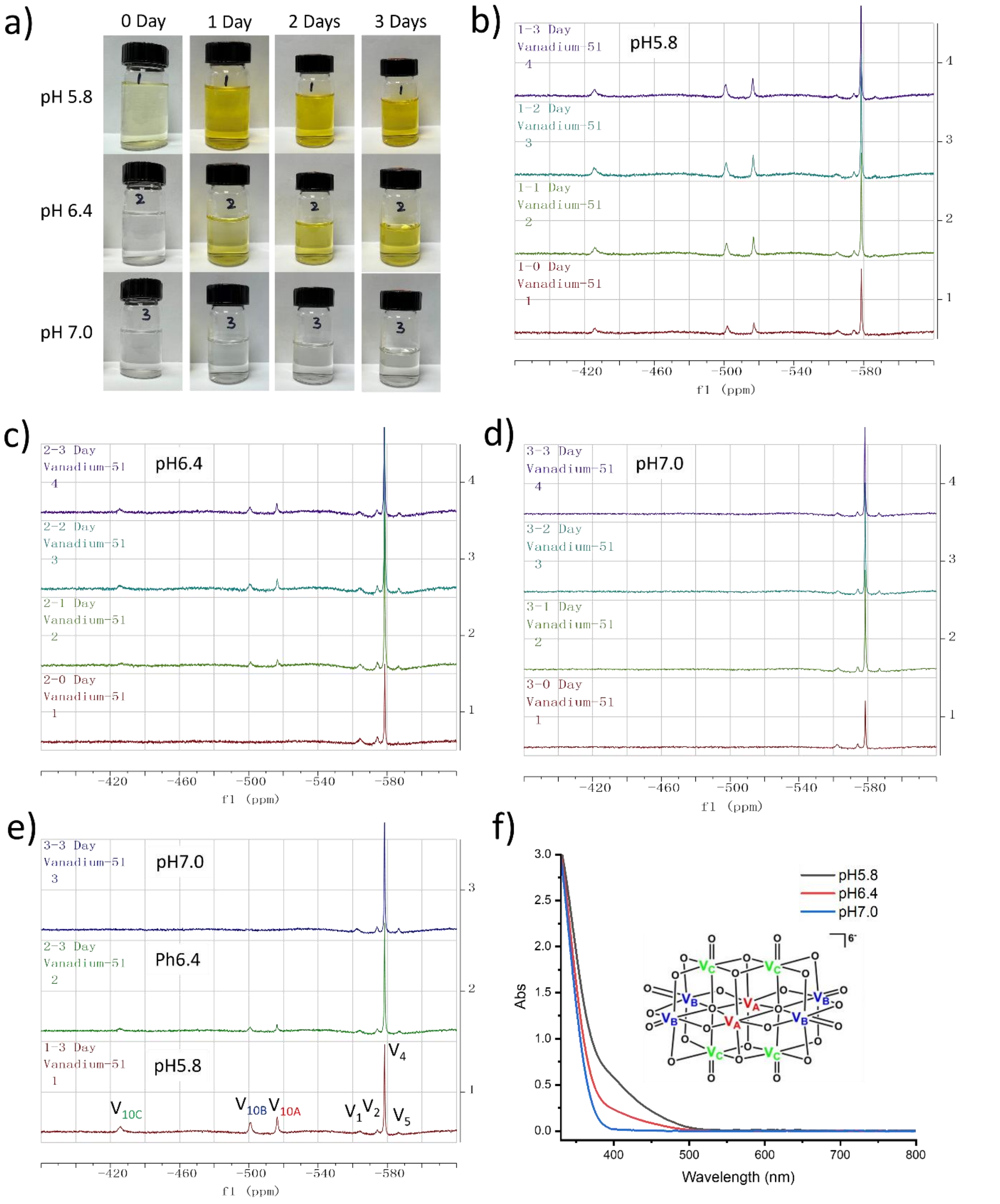


**Fig. S4 | Formation of polyoxovanadates from $NH_4VO_3$ under controlled pH conditions.** (**a**) Vanadate solutions prepared in 1× sodium phosphate buffer at pH 5.8, 6.4, and 7.0 separately. (**b–e**) $^{51}V$ NMR spectra of vanadate species as a function of pH and incubation time. (**f**) UV–vis absorption spectra of vanadate solutions at different pH values after 3 days of incubation; inset shows the crystal structure of the $V_{10}$ cluster. Reproduced with permission from Elsevier.[15]

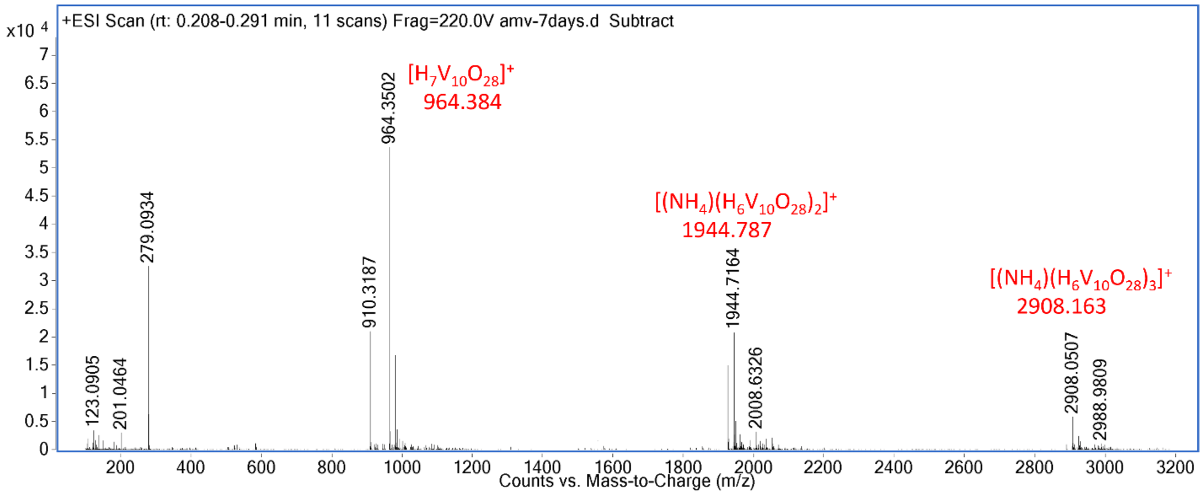


**Fig. S5 | ESI–TOF mass spectrometric analysis of vanadate species at pH 5.8**. Electrospray ionization time-of-flight (ESI–TOF) mass spectrum was recorded in positive-ion mode. The mobile phase consisted of 10% $H_2O$ and 90% acetonitrile at a flow rate of 1.0 mL min$^{-1}$. Values shown beneath the molecular formulae correspond to calculated exact masses for the assigned species.

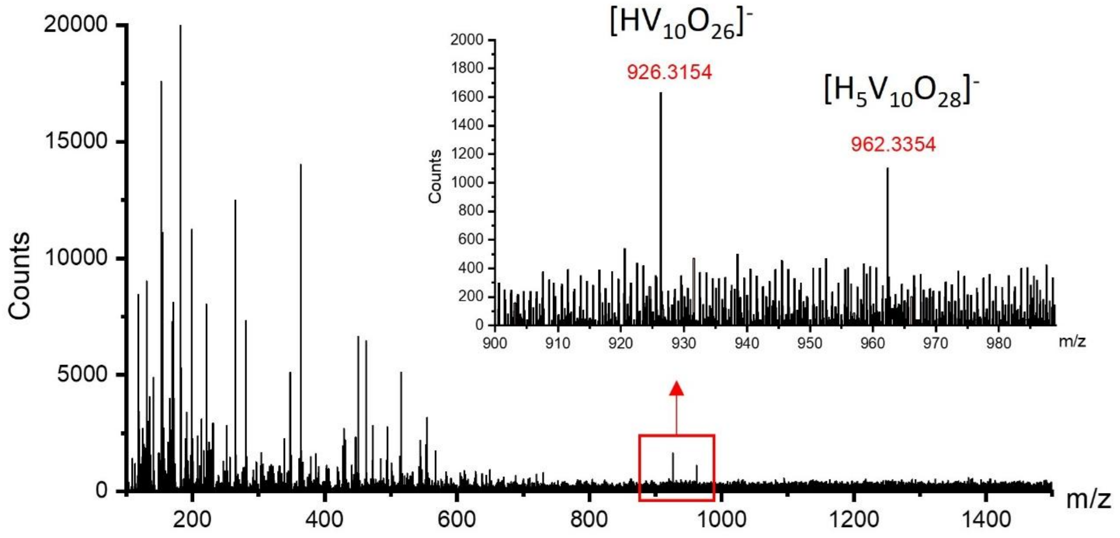


**Fig. S6 | ESI–TOF mass spectrometric analysis of *L*-oxovanadate clusters in negative-ion mode**. Calculated exact masses for the assigned species are 926.347 Da for $[HV_{10}O_{26}]^-$ and 962.368 Da for $[H_5V_{10}O_{28}]^-$.

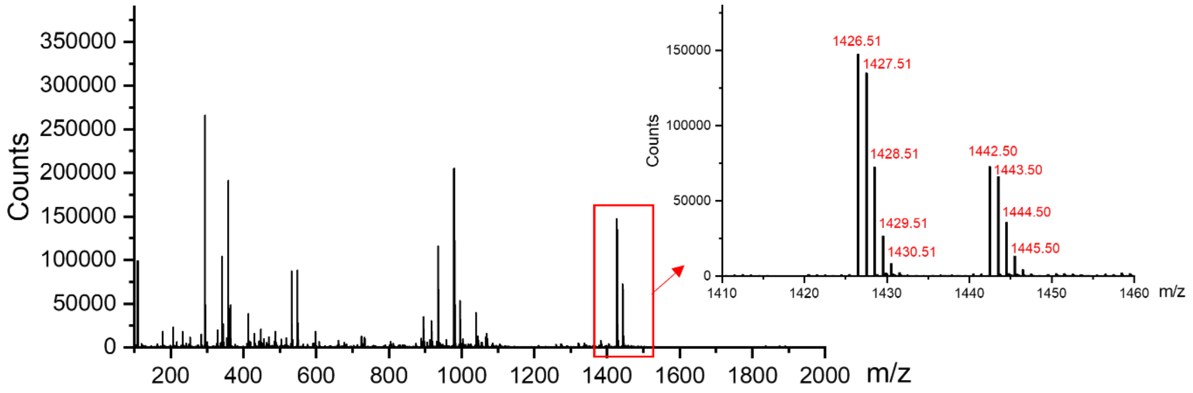


**Isotopic Abundances for**

$H_5V_{10}O_{28}(C_4H_4O_4)_4$

| Mass/Charge | Fraction | Intensity |
|---|---|---|
| 1423.41219 | 0.0000014 | 0.00 |
| 1424.41219 | 0.0002081 | 0.03 |
| 1425.41219 | 0.0184711 | 2.50 |
| 1426.41219 | 0.7389681 | 100.00 |
| 1427.41219 | 0.1438990 | 19.47 |
| 1428.41219 | 0.0798384 | 10.80 |
| 1429.41219 | 0.0136629 | 1.85 |
| 1430.41219 | 0.0041557 | 0.56 |
| 1431.41219 | 0.0006372 | 0.09 |
| 1432.41219 | 0.0001393 | 0.02 |
| 1433.41219 | 0.0000194 | 0.00 |
| 1434.41219 | 0.0000034 | 0.00 |

**Isotopic Abundances for**

$H_5V_{10}O_{28}(C_4H_4O_4)_3(C_4H_4O_5)$

| Mass/Charge | Fraction | Intensity |
|---|---|---|
| 1439.40710 | 0.0000014 | 0.00 |
| 1440.40710 | 0.0002076 | 0.03 |
| 1441.40710 | 0.0184263 | 2.50 |
| 1442.40710 | 0.7371799 | 100.00 |
| 1443.40710 | 0.1438680 | 19.52 |
| 1444.40710 | 0.0812140 | 11.02 |
| 1445.40710 | 0.0139551 | 1.89 |
| 1446.40710 | 0.0043145 | 0.59 |
| 1447.40710 | 0.0006653 | 0.09 |
| 1448.40710 | 0.0001477 | 0.02 |
| 1449.40710 | 0.0000207 | 0.00 |
| 1450.40710 | 0.0000037 | 0.00 |

**Fig. S7 | ESI–TOF mass spectrometric characterization of *L*-oxovanadate clusters**. ESI–TOF mass spectrum of *L*-oxovanadate recorded in positive-ion mode, together with calculated isotopic distributions for $[H_5V_{10}O_{28}(C_4H_4O_4)_4]^+$ and $[H_5V_{10}O_{29}(C_4H_4O_4)_4]^+$ shown at the bottom.

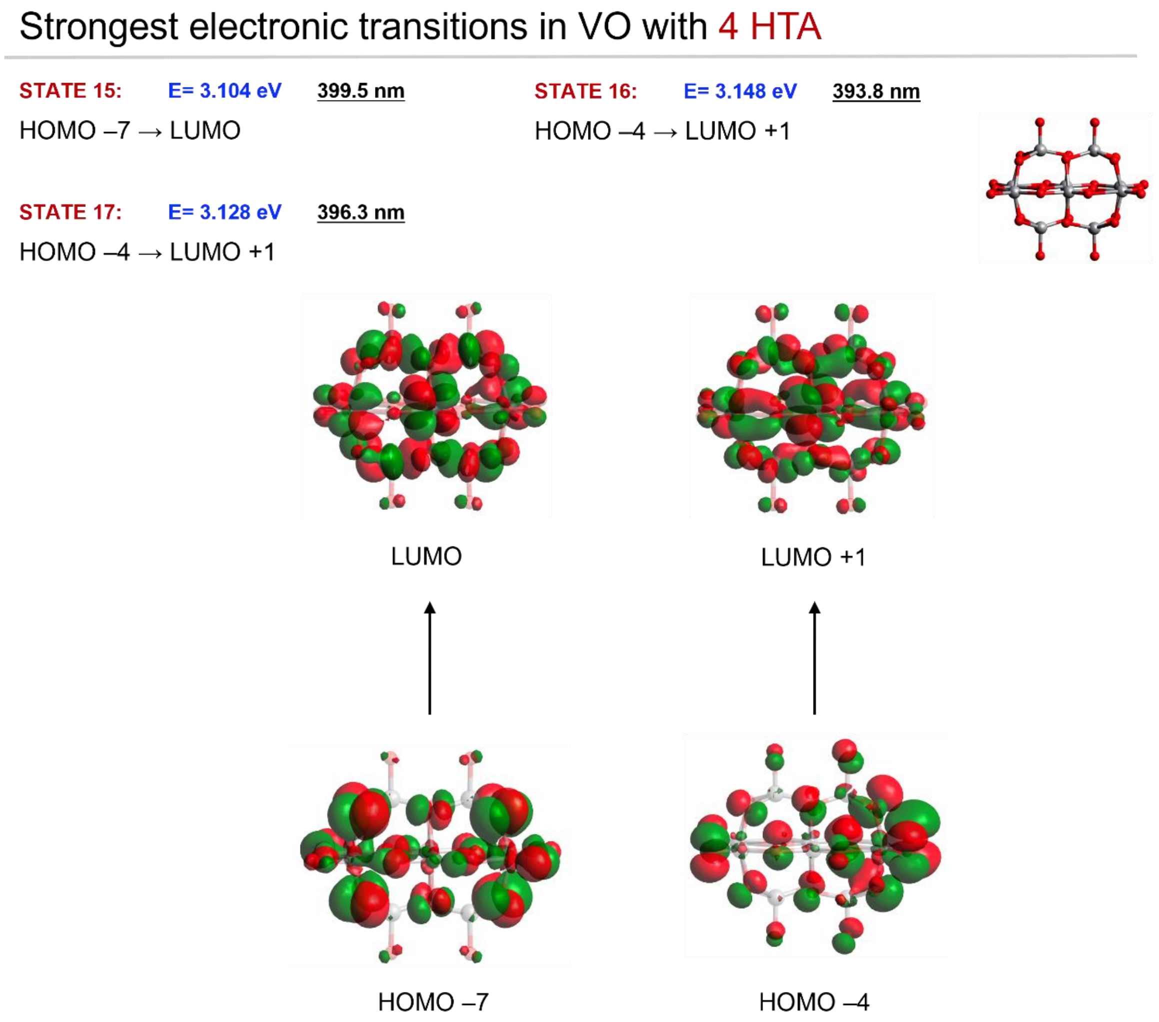


**Fig. S8 | Calculated electronic transitions underlying the dominant CD features of $V_{10}O_{28}(HTA)_4$.**

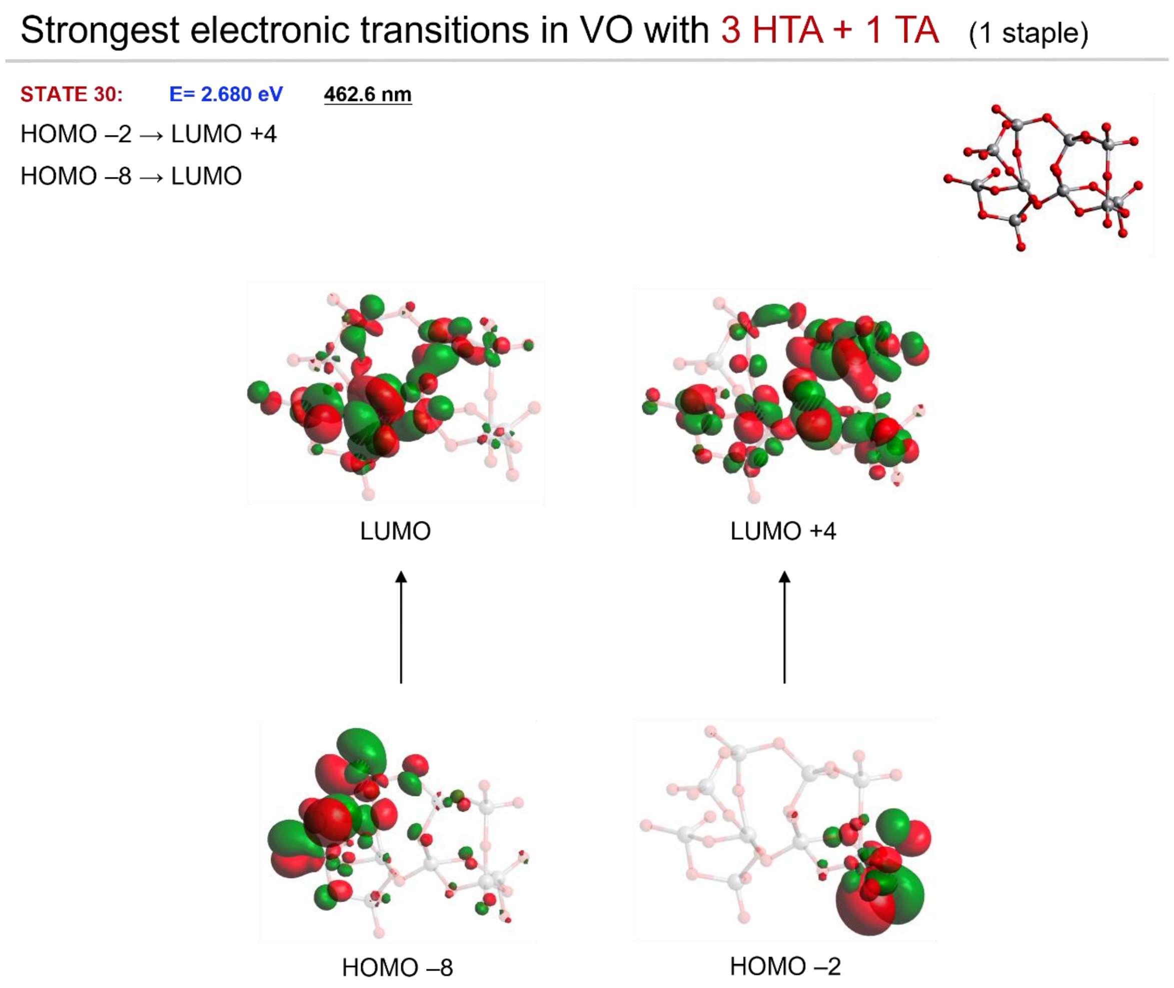


**Fig. S9 | Calculated electronic transitions underlying the dominant CD features of $V_{10}O_{28}(TA)(HTA)_3$.**

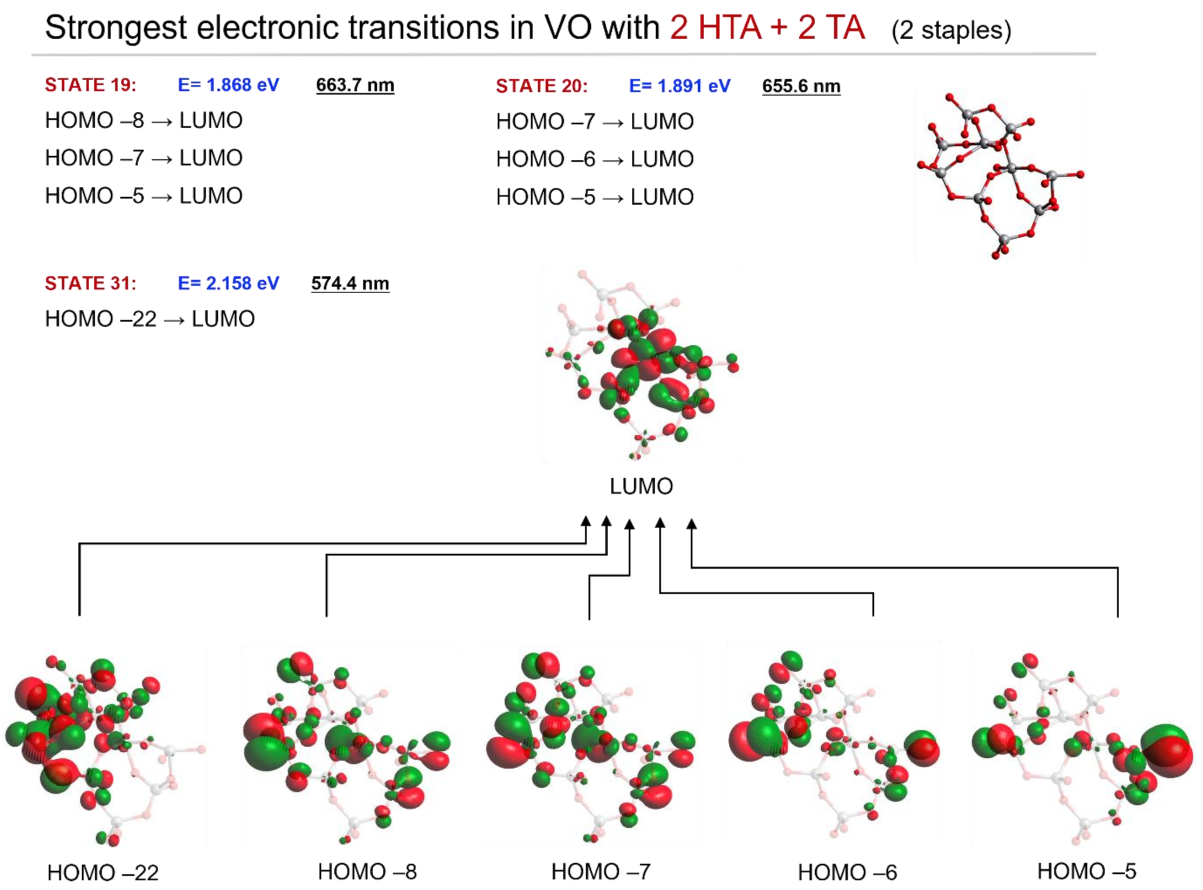


**Fig. S10 | Calculated electronic transitions underlying the dominant CD features of $V_{10}O_{28}(TA)_2(HTA)_2$.**

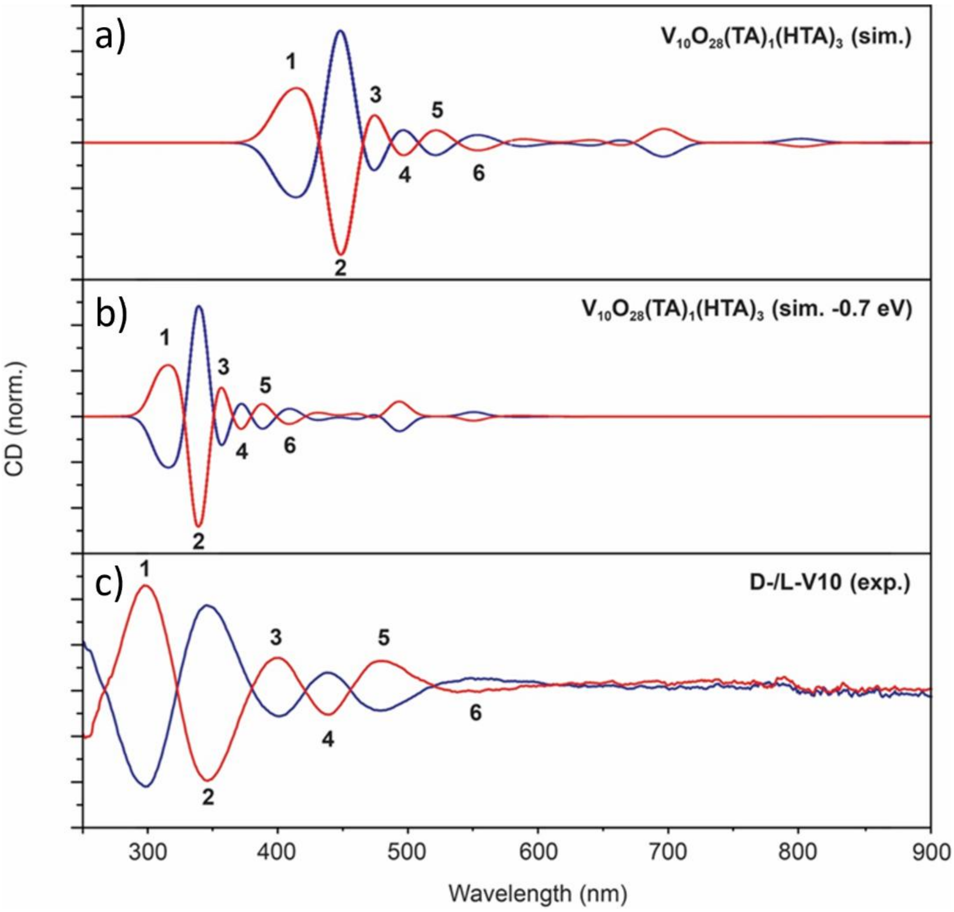


**Fig. S11 | Comparison of simulated and experimental CD spectra of chiral $V_{10}(TA)_4$.** (**a**) Simulated CD spectra of *L*- and *D*-$V_{10}(TA)_4$. (**b**) Simulated spectra from panel a after application of a 0.7 eV blue shift to align peak positions with the experimental spectra. (**c**) Experimental CD spectra of *L*- and *D*-$V_{10}(TA)_4$ recorded in acetate buffer at pH 4; the *L*-form is shown in blue and the *D*-form in red.

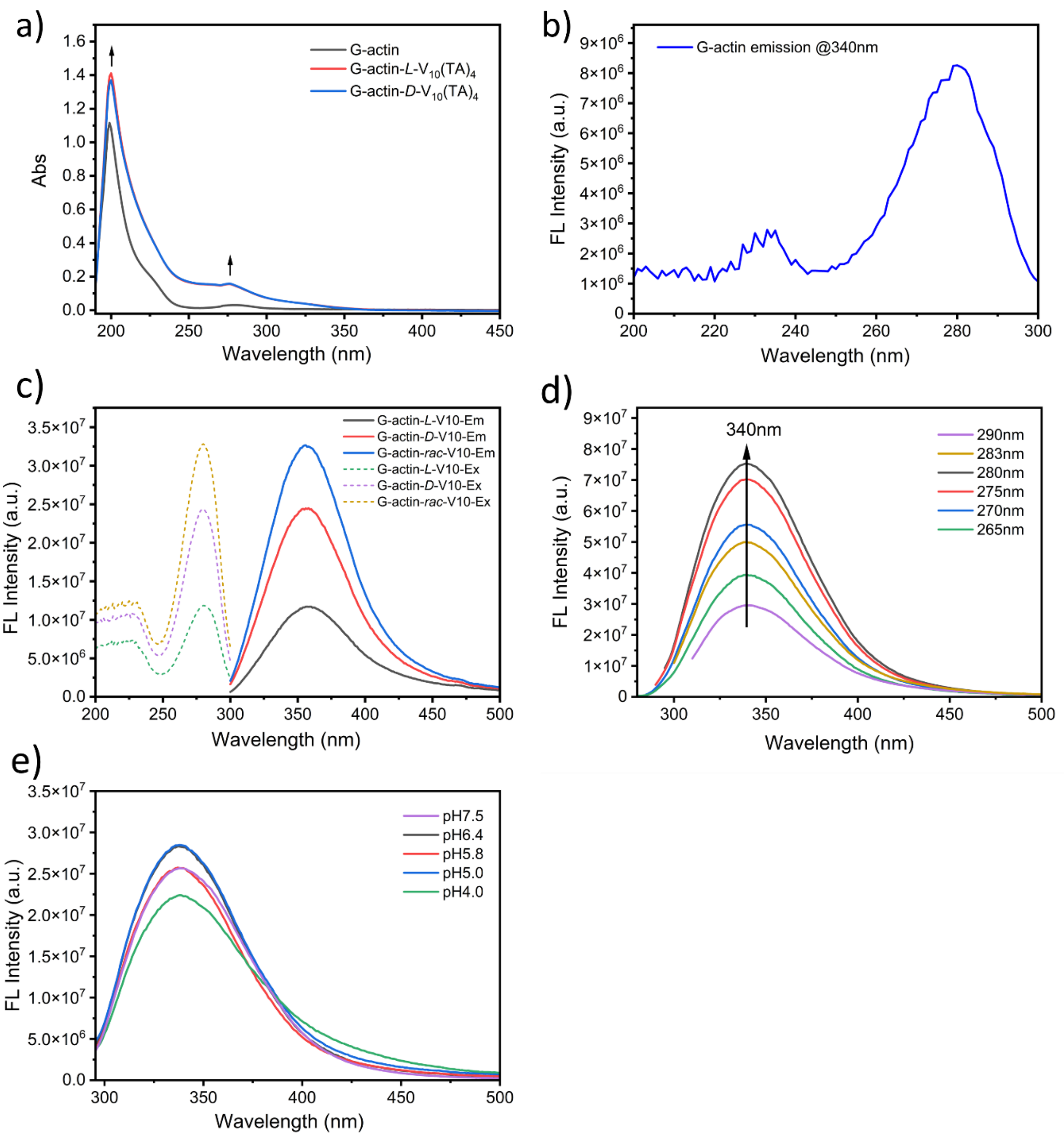


**Fig. S12 | Absorption and fluorescence characterization of G-actin in the presence of $V_{10}(TA)_4$.** (**a**) UV–vis absorption spectra of 2 μM G-actin and mixtures of G-actin with *L*- and *D*-$V_{10}(TA)_4$. (**b**) Excitation spectrum of G-actin (λ_ex = 280 nm) monitored at an emission wavelength of 340 nm. (**c**) Emission (solid lines) and excitation (dashed lines) spectra of mixtures containing 2 μM G-actin and 160 μM *L*-, *D*-, or *rac*-$V_{10}(TA)_4$. (**d**) Emission spectra of 2 μM G-actin recorded under different excitation wavelengths. (**e**) Emission spectra of 2 μM G-actin recorded at λ_ex = 280 nm under varying pH conditions.

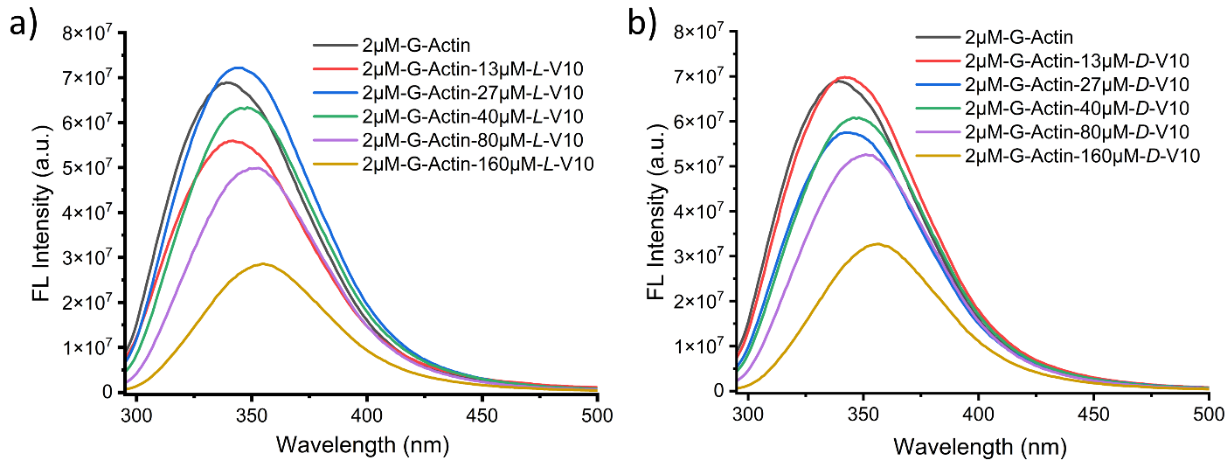


**Fig. S13 | Fluorescence emission spectra of G-actin in the presence of chiral $V_{10}(TA)_4$.** Fluorescence emission spectra of 2 μM G-actin in G-buffer recorded in the presence of increasing concentrations of *L*-$V_{10}(TA)_4$ (**a**) and *D*-$V_{10}(TA)_4$ (**b**), with an excitation wavelength of 280 nm.

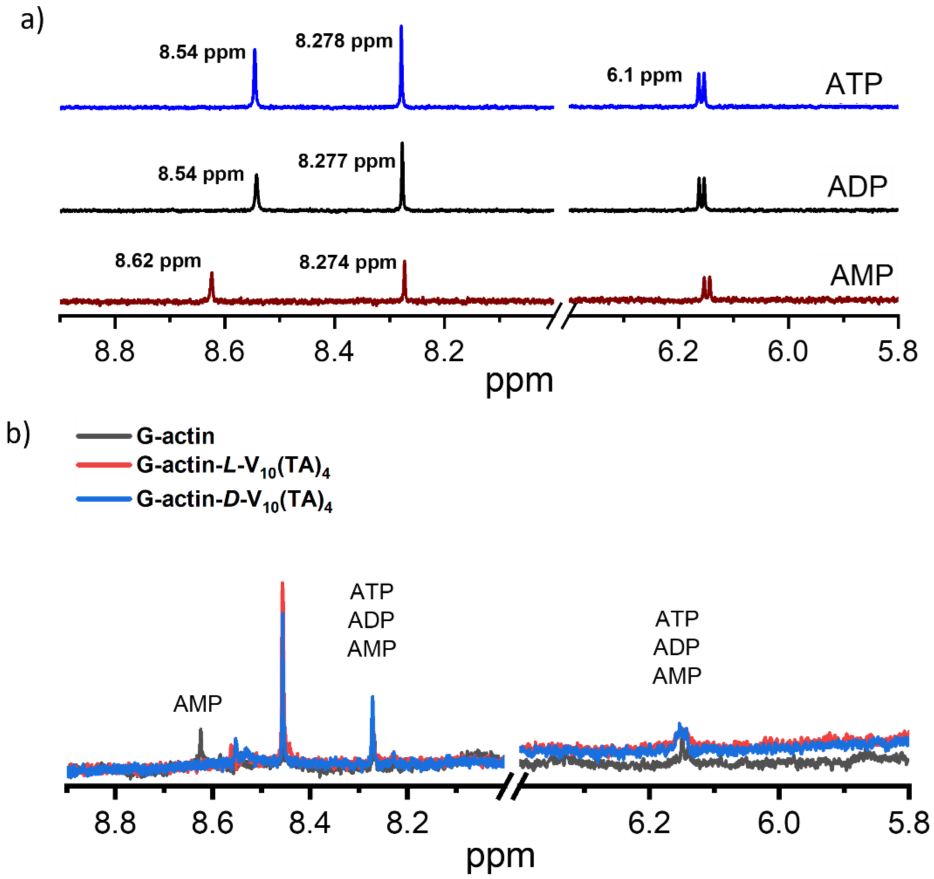


**Fig. S14 | $^{1}$H NMR characterization of nucleotide species and actin–$V_{10}(TA)_4$ interactions.** (**a**) $^{1}$H NMR spectra of ATP, ADP, and AMP separately. The resonance at 8.62 ppm is characteristic of AMP. (**b**) $^{1}$H NMR spectra of G-actin (black), G-actin-*L*-$V_{10}(TA)_4$ (red line) and G-actin-*D*-$V_{10}(TA)_4$ (blue line).

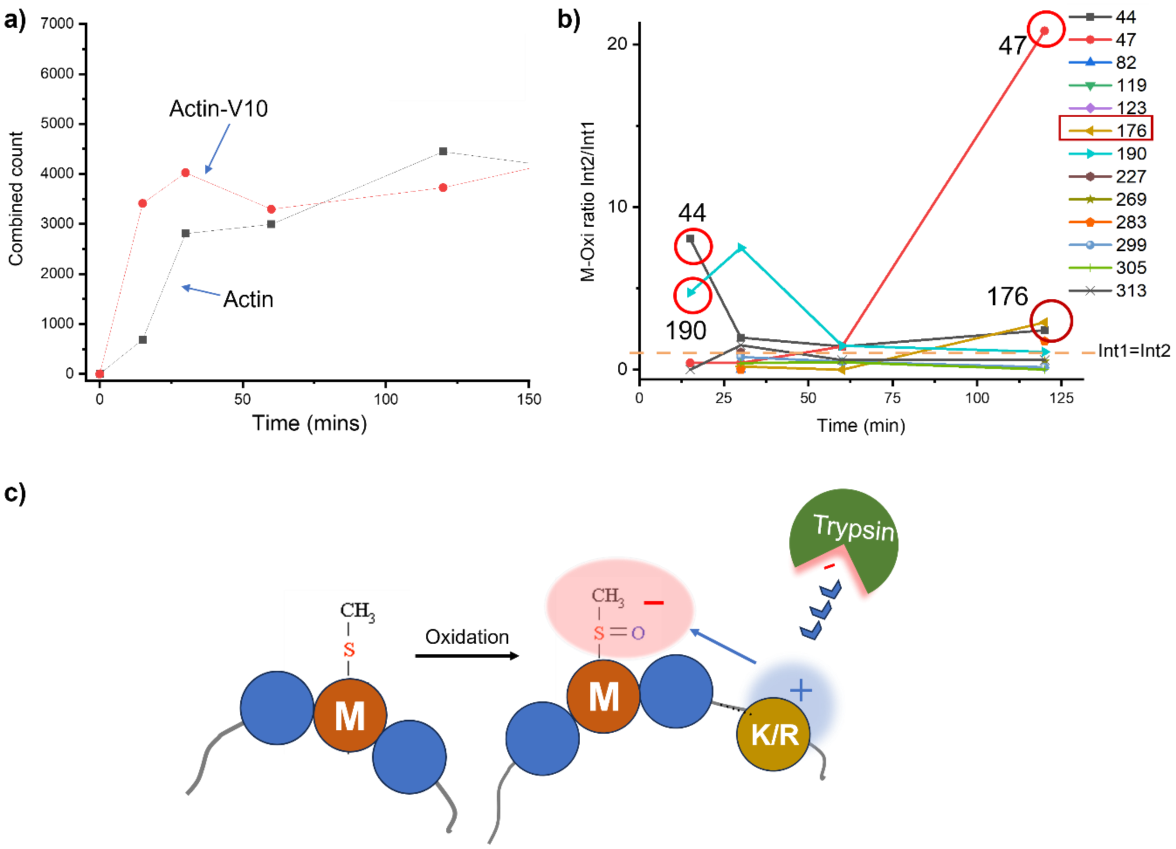


**Fig. S15 | Kinetic analysis of trypsin digestion of actin by LC–MS/MS**. (**a**) Trypsin digestion kinetics of actin quantified from combined peptide intensities obtained by LC–MS/MS analysis. The red curve corresponds to the actin–*L*-$V_{10}(TA)_4$ system, and the black curve corresponds to actin alone. (**b**) Time-dependent methionine oxidation index for 14 identified residues within G-actin. (**c**) Schematic illustrating the proposed mechanism underlying accelerated trypsin digestion of G-actin following Methionine oxidation.

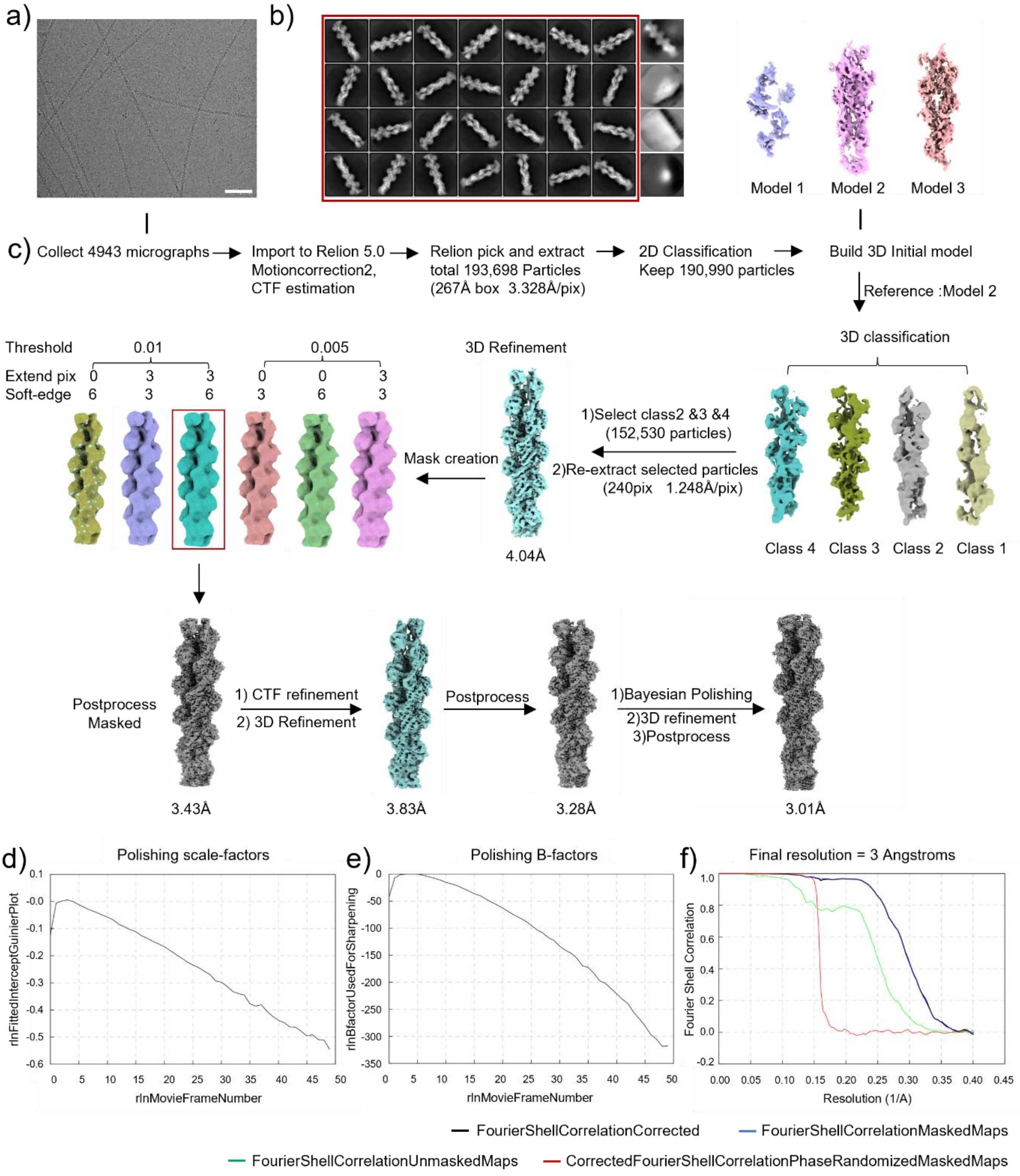


**Fig. S16 | Cryo-EM image processing workflow for F-actin formed in the actin–*D*-$V_{10}(TA)_4$ system.** (**a**) Representative cryo-EM micrograph from a dataset comprising 4,943 collected micrographs. (**b**) Two-dimensional class averages of F-actin particles generated using RELION 5.0; classes selected for further processing are indicated by red boxes, while discarded classes are shown without highlighting. The box size is 267 × 267 Å². (**c**) Image-processing workflow used for F-actin reconstruction. All steps were performed in RELION 5.0; following 3D classification, particles are displayed in a common orientation. (**d**) Bayesian polishing scale-factor plot. (**e**) Bayesian polishing B-factor plot. (**f**) Fourier shell correlation (FSC) curves of the F-actin reconstruction before and after post-processing, calculated in RELION 5.0.

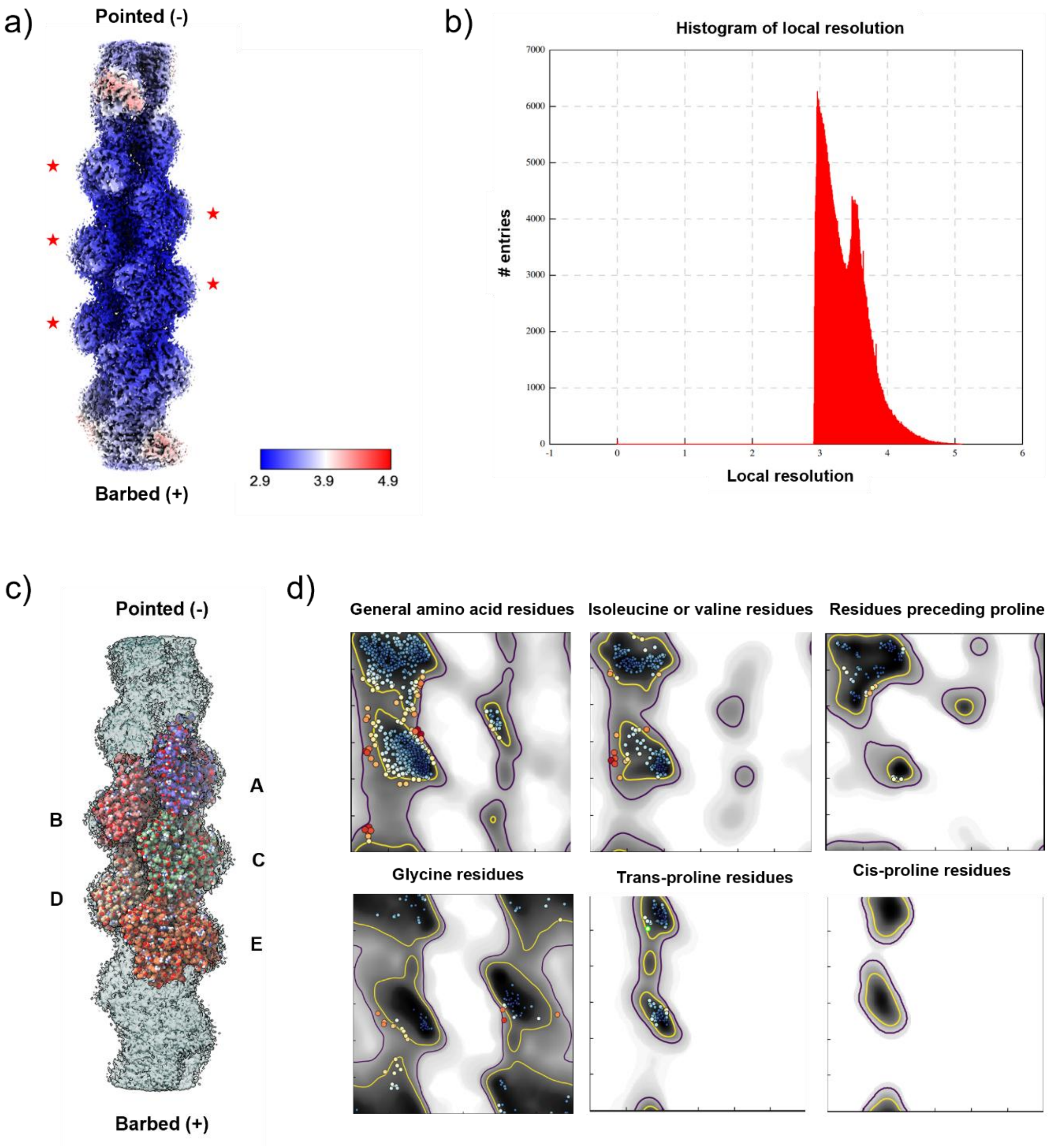


**Fig. S17 | Local resolution analysis and model validation of the reconstructed F-actin structure.** (**a**) Local resolution map (Å) of the reconstructed F-actin density. Density corresponding to five labeled actin subunits was selected for subsequent coordinate model refinement. (**b**) Overall histogram of local resolution values generated using RELION. (**c**) Docking of the atomic model (PDB: 8A2Y) into the reconstructed 3.0 Å F-actin density map; for clarity, the density map is rotated 180° about the z axis relative to panel a. (**d**) Ramachandran plot of the refined coordinate model (PDB: 000010LU) following ISOLDE refinement.

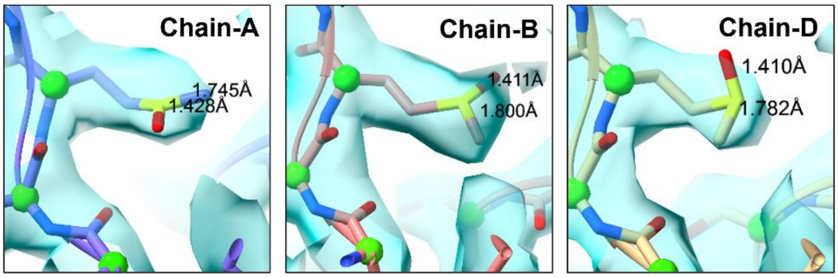


**Fig. S18 | Locally modified Met-178 side chain to Methionine sulfoxide.** Modified Met-176 residues in chain A, chain B and Chain D by manually introducing an extra oxygen atom to the sulfur and aligned with the density map.

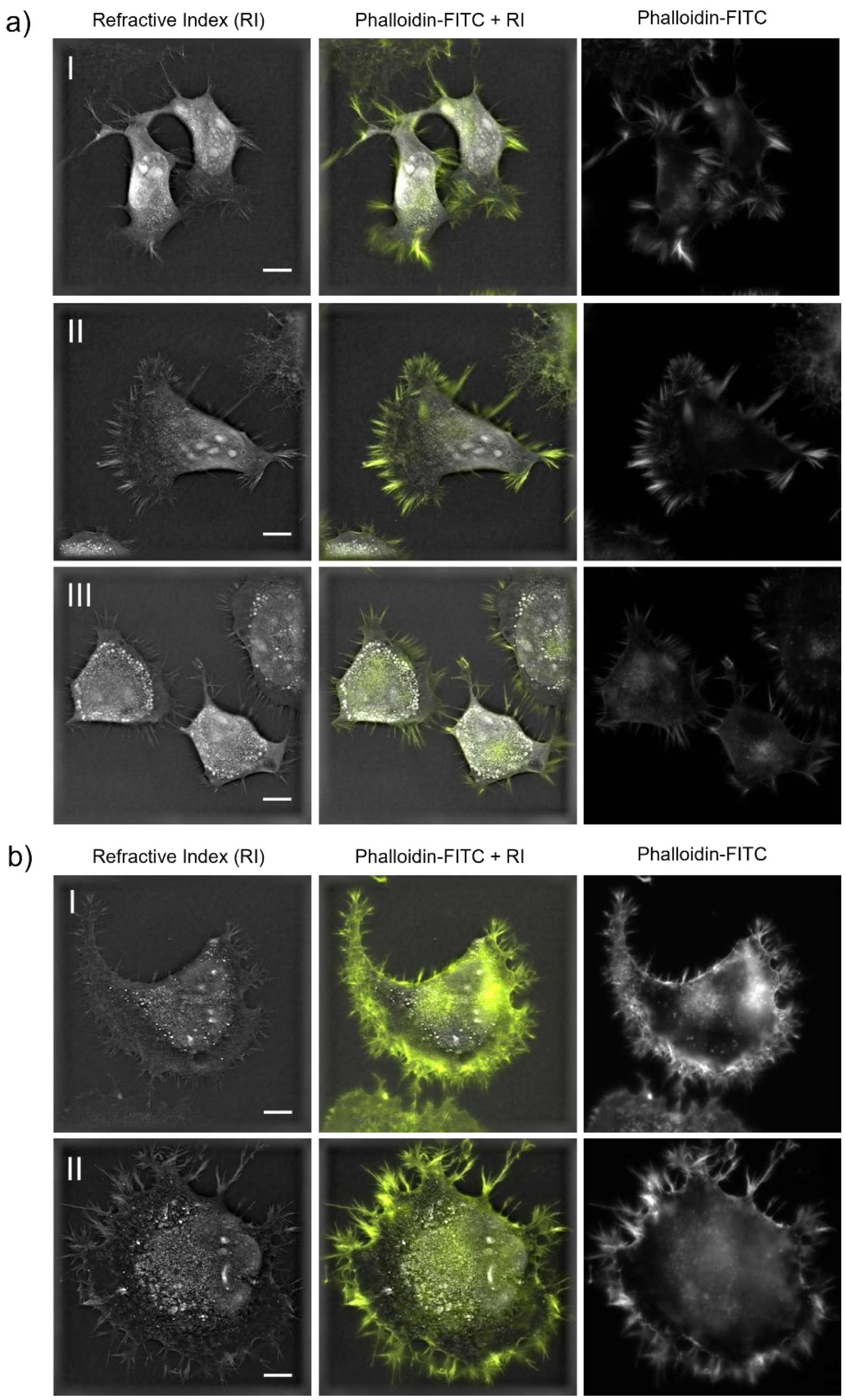


**Fig. S19 | Colocalization of F-actin inside NG108-15 cells with Refractive index and Phalloidin-FITC fluorescence visualized under Nanolive 3D cell explorer.** (a) Three cases of initial NG108-15 cells and (b) two cases of NG108-15 cells been incubated with 5 μM *D*-$V_{10}(TA)_4$. Phalloidin-FITC (200nM stain concentration) was applied after cell been fixed with 4% Paraformaldehyde and permeabilizing with 0.1% Triton X-100, scale bar 10 μm.

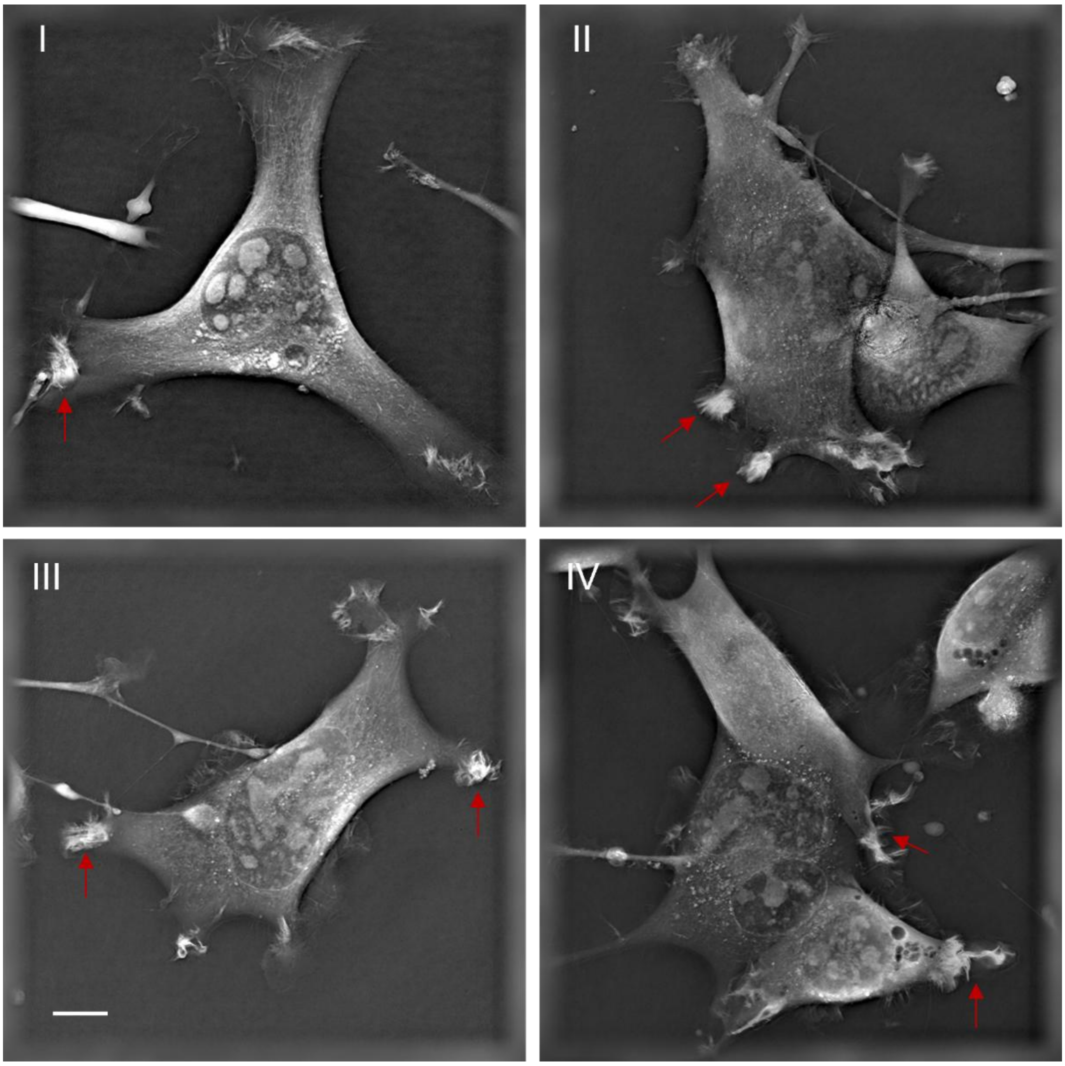


**Fig. S20 | Representation of NG108-15 cells co-incubated with 7 μM *L*-$V_{10}$(TA)$_4$.** Red arrows indicated the F-actin accumulation at the growth cone after co-incubation with chiral decavanadate nanoclusters for more than 5h, images were recorded by the Nanolive 3D cell explorer, scale bar 10 μm.

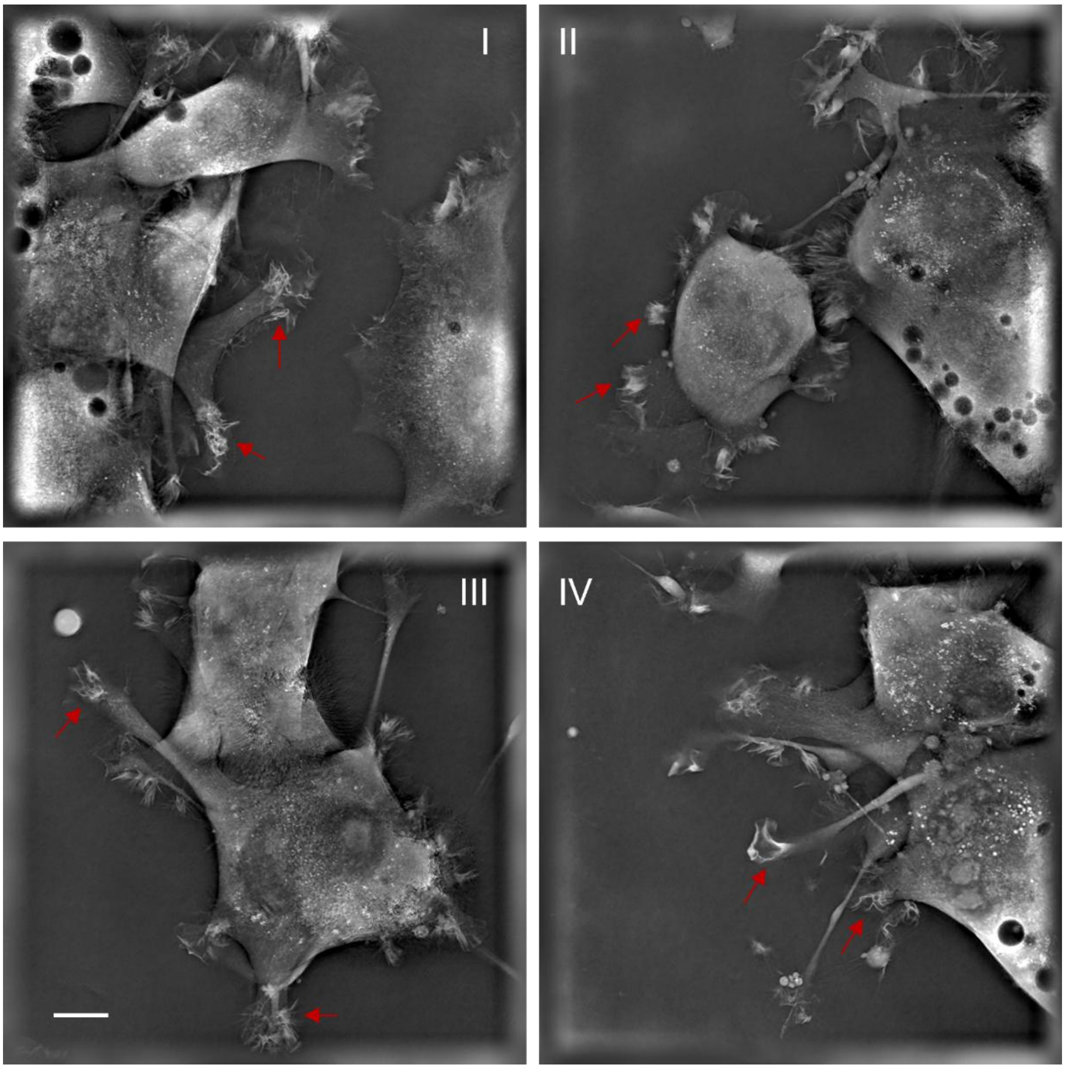


**Fig. S21 | Representation of NG108-15 cells co-incubated with 7 μM *D*-$V_{10}(TA)_4$.** Red arrows indicated the F-actin accumulation at the growth cone after co-incubation with chiral decavanadate nanoclusters for more than 5h, images were recorded by the Nanolive 3D cell explorer, scale bar 10 μm.

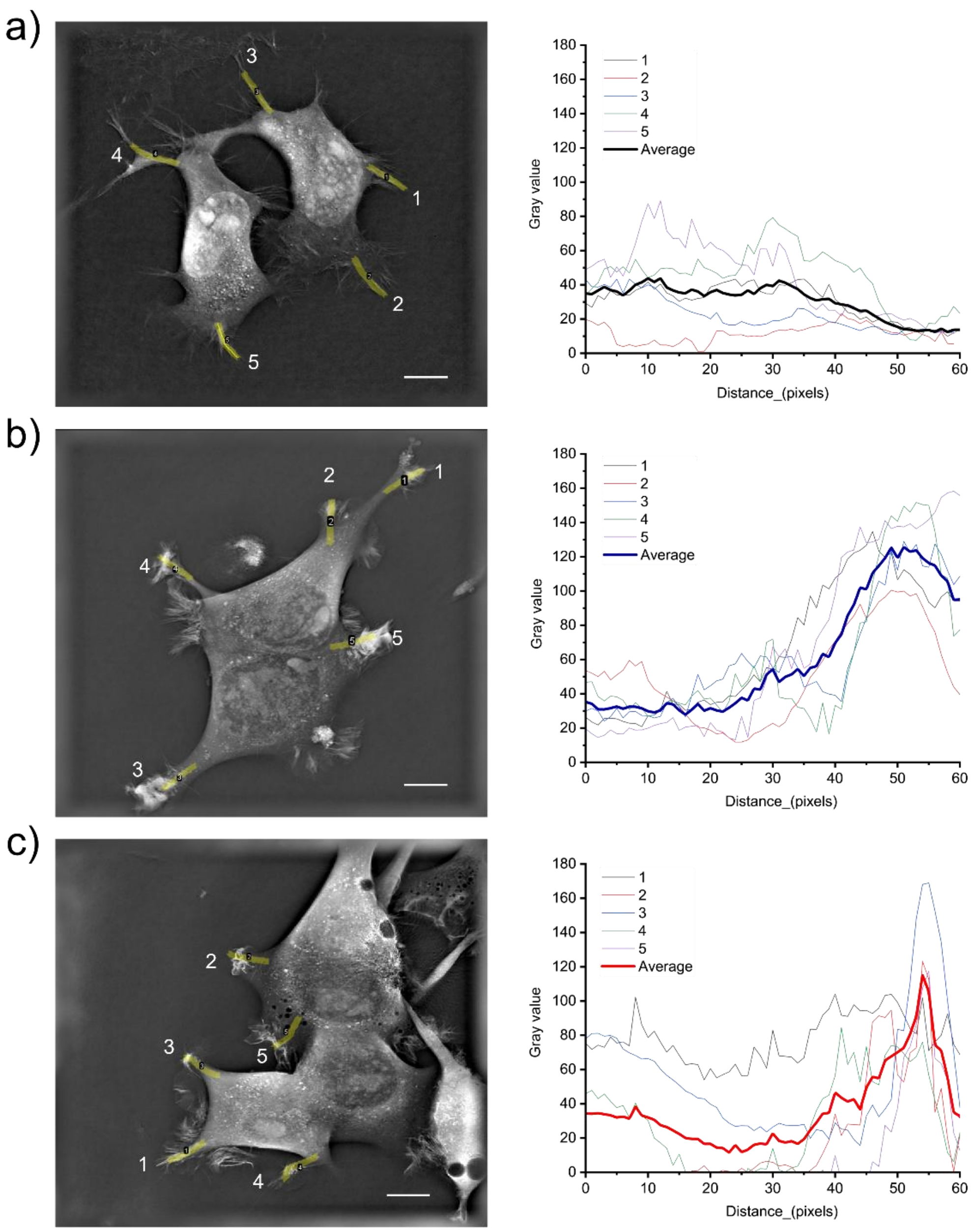


**Fig. S22 | Mass intensity distribution along neurite shaft to growth cone in various NG108-15 cell condition.** Selected 12μm segments from neurite shaft to growth cone tip with in (a) original NG108-15 cells, (b) NG108-15-*L*-$V_{10}(TA)_4$ cells, and (c) NG108-15-*D*-$V_{10}(TA)_4$ cells. Nanolive 3D cell explorer recorded micrographs were displayed in 512 x 512 pixels with real size 103 μm x103 μm, scale bar 10μm.

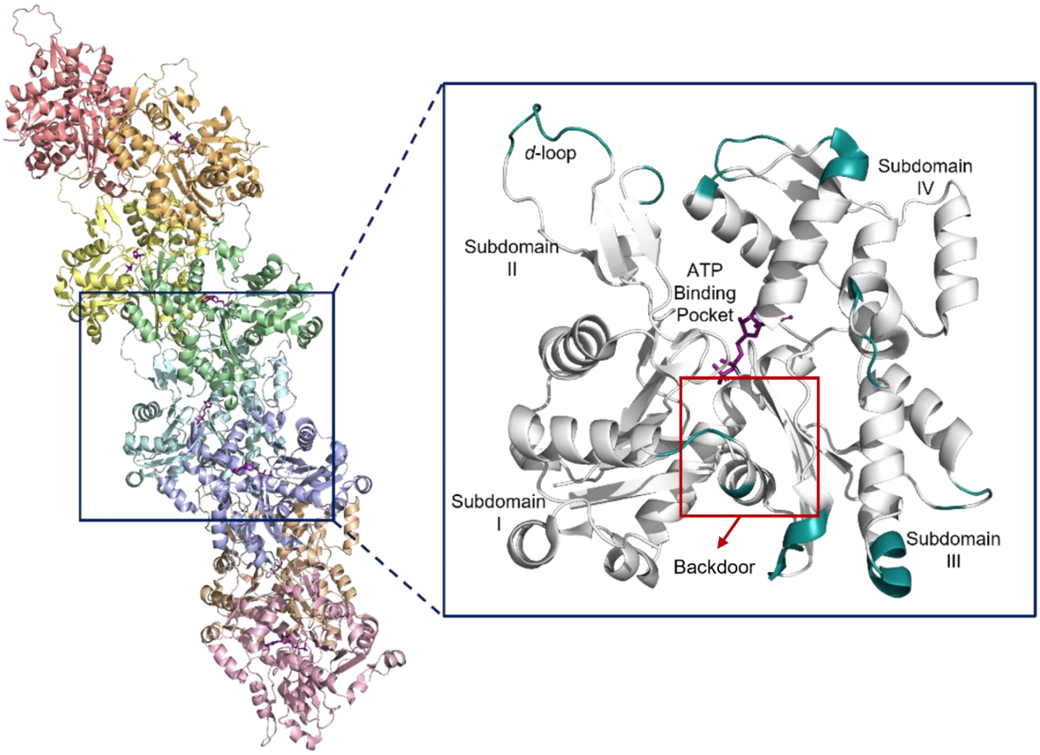


**Fig. S23 | Structural context for actin filament contacts and the Pi-release backdoor**. Structure of the F-actin filament from *Oryctolagus cuniculus* (PDB ID: 6BNO).[16] Individual actin subunits are shown in different colors. The boxed region indicates the location of the representative G-actin monomer (AlphaFold Protein Structure P68135) shown at higher magnification (right). In the enlarged view, residues involved in inter-subunit contacts are highlighted, and ADP is shown in purple. The region associated with inorganic phosphate (Pi) release ("backdoor") following ATP hydrolysis is indicated.

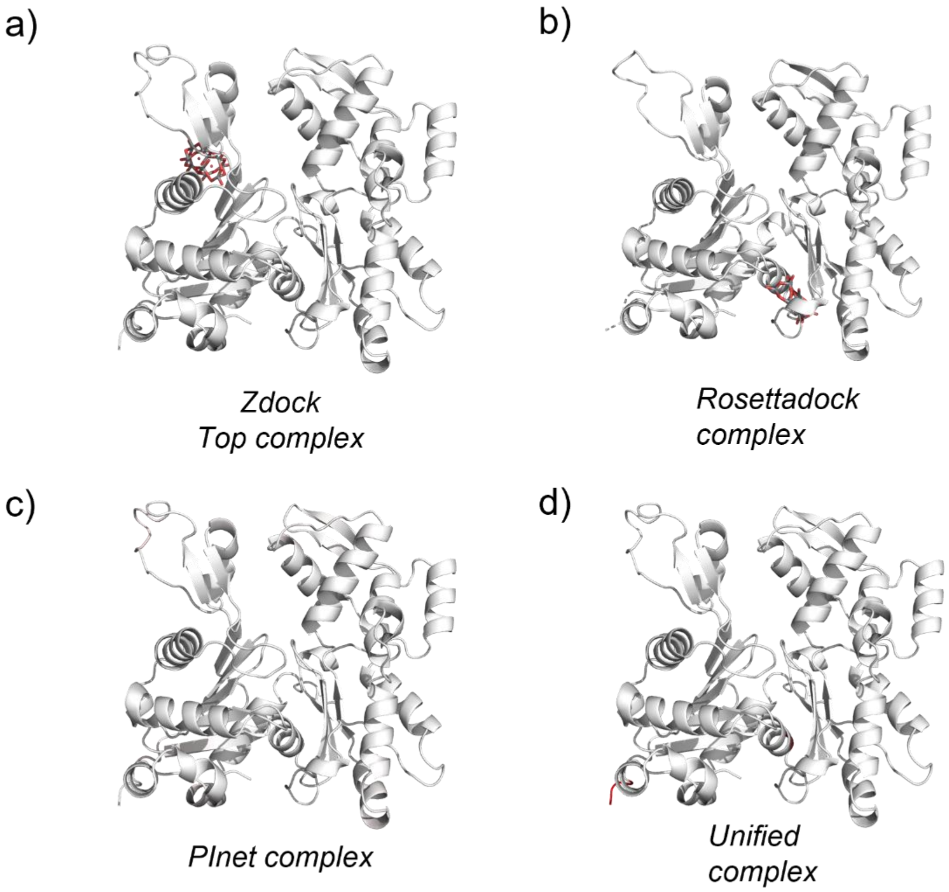


**Fig. S24 | Prediction of actin–$V_{10}O_{28}$ nanocluster complex by four models**. Individual panels show representative docking poses for single nanocluster displayed in the same molecular orientation.

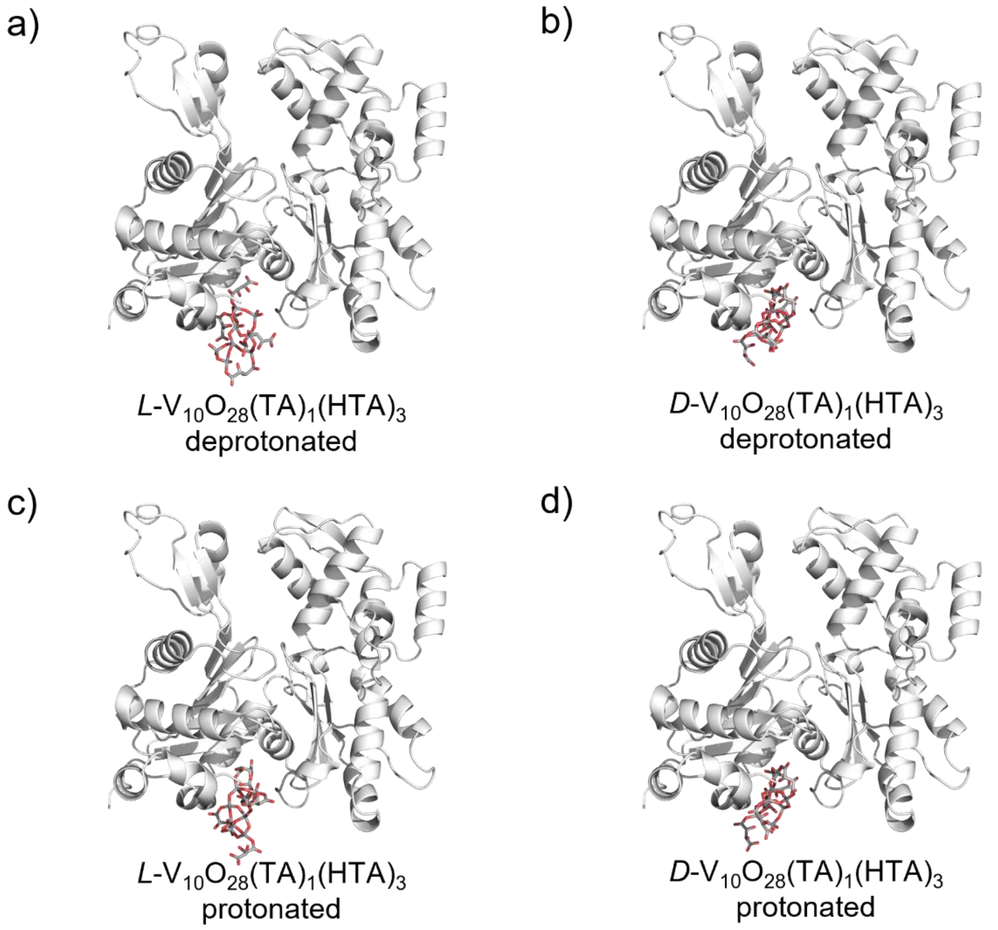


**Fig. S25 | *ZDOCK* prediction of actin–*L/D*-$V_{10}O_{28}(TA)_1(HTA)_3$ nanocluster docking**. Rigid-body docking predictions between G-actin and individual nanoclusters with different protonation states, and chirality, calculated using *ZDOCK*. Individual panels show representative docking poses for single nanoclusters displayed in the same molecular orientation.

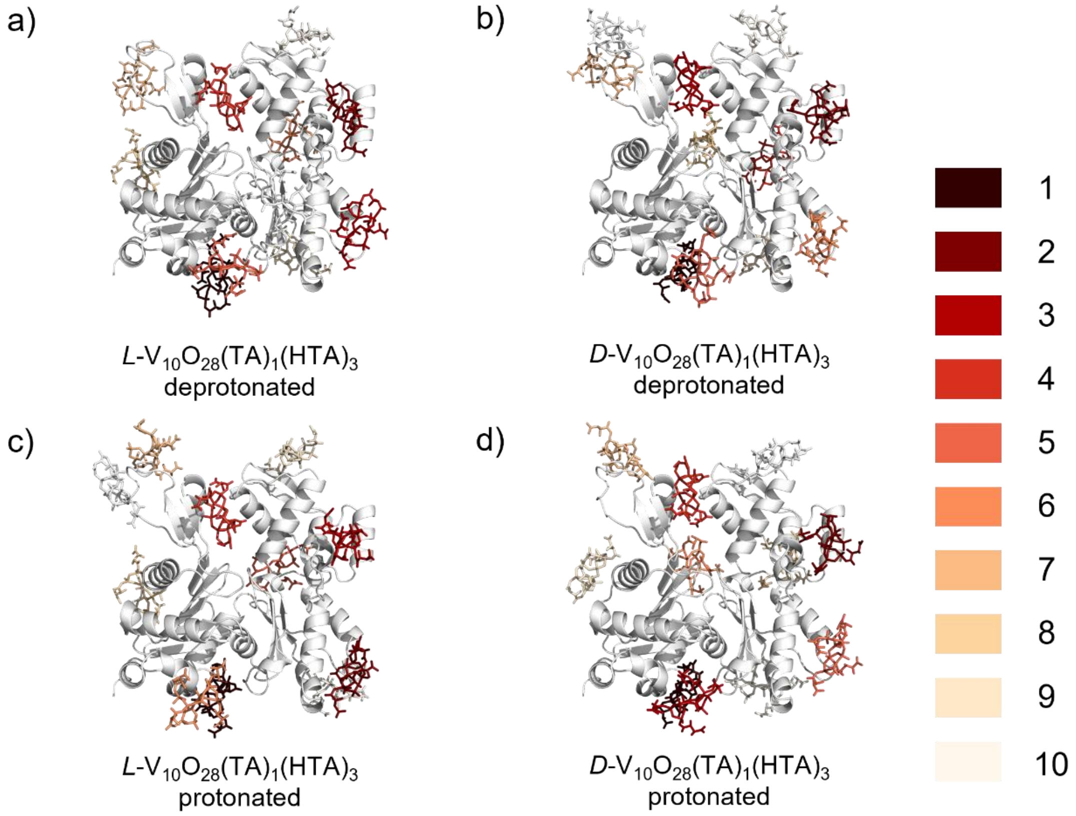


**Fig. S26 | *ZDOCK* prediction of actin–*L/D*-$V_{10}O_{28}(TA)_1(HTA)_3$ docking with multiple nanoclusters**. Predicted rigid-body docking of G-actin with up to ten nanoclusters of different protonation states, and chirality, calculated using *ZDOCK*. Individual panels show representative docking poses for multiple nanoclusters displayed in the same molecular orientation. Nanoclusters are colored according to discrete *ZDOCK* docking score ranges, from lower scores (lighter colors) to higher scores (dark red), as indicated by the color legend.

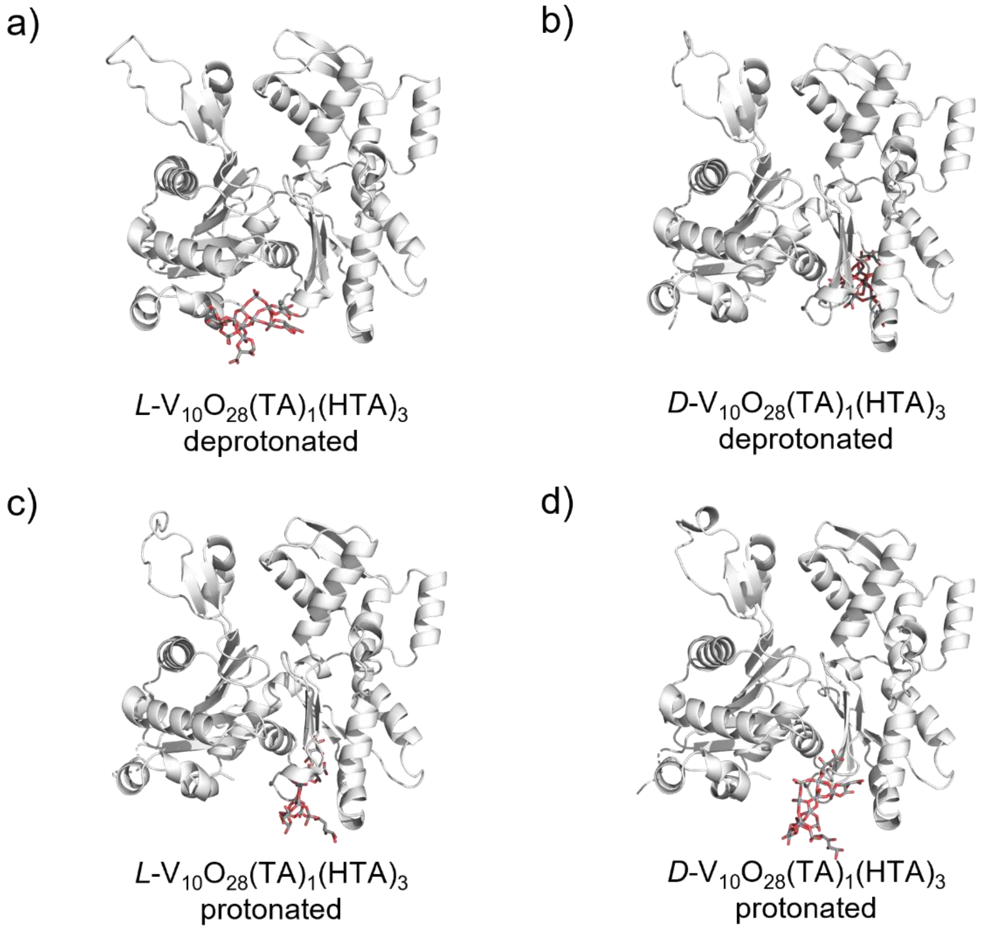


**Fig. S27 | *RosettaFold All-Atom* prediction of actin–*L/D*-$V_{10}O_{28}(TA)_1(HTA)_3$ nanocluster docking**. Predicted docking between G-actin and single nanoclusters with different protonation states, and chirality, calculated using *RosettaFold All-Atom* (RFAA). Individual panels show representative docking poses displayed in the same molecular orientation.

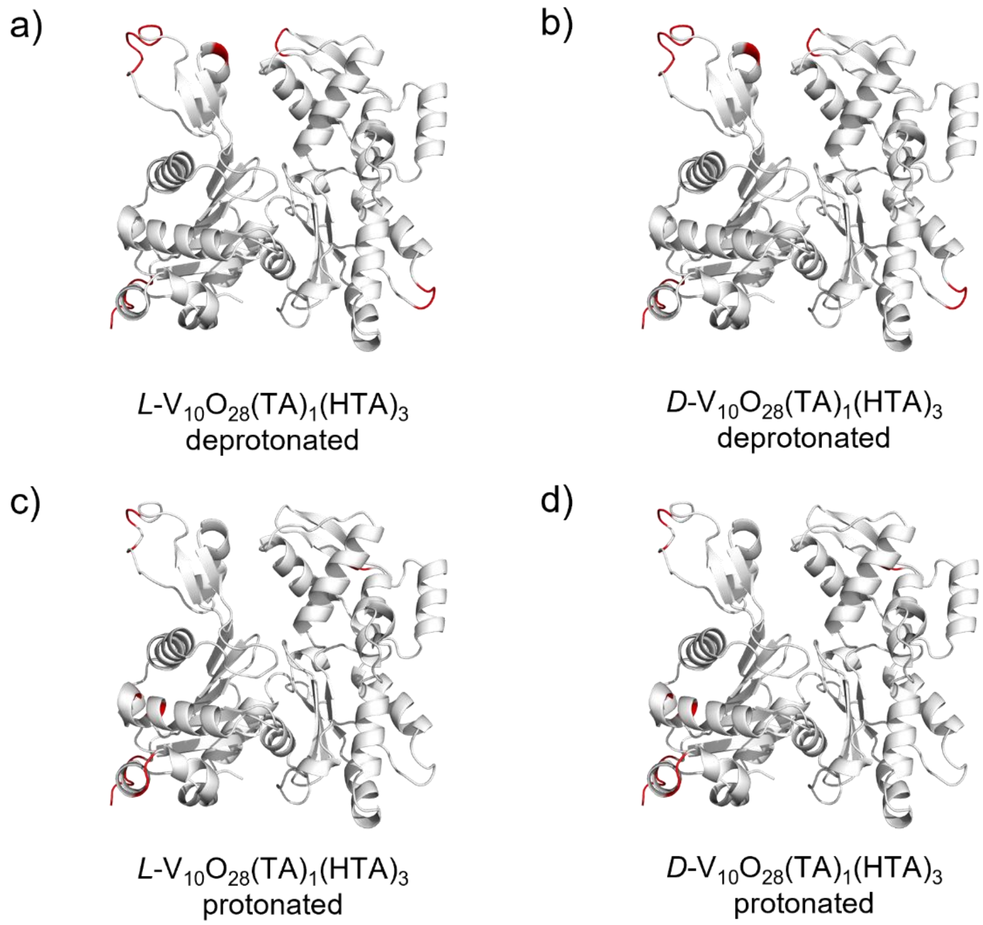


**Fig. S28 | *Unified* model prediction of actin regions involved in *L/D*-$V_{10}O_{28}(TA)_1(HTA)_3$ nanocluster docking.** Predicted interaction regions between G-actin and single nanoclusters with different protonation states, and chirality, calculated using the *Unified* model. Individual panels show G-actin displayed in the same molecular orientation, with residues predicted to participate in nanocluster docking highlighted.

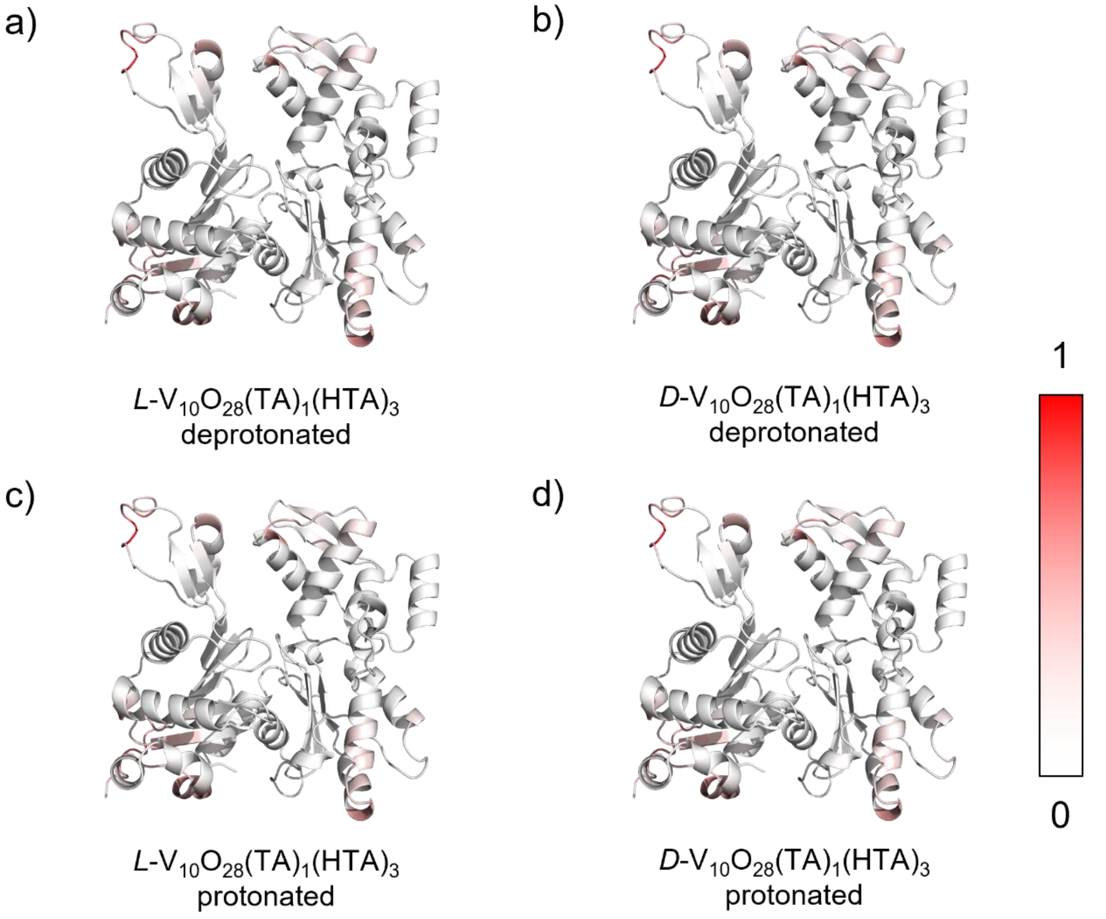


**Fig. S29 | *PInet* prediction of actin–*L/D*-$V_{10}O_{28}(TA)_1(HTA)_3$ nanocluster docking efficiency**. Prediction of docking efficiency between G-actin and nanoclusters with different protonation states, and chirality, calculated using the *Protein Interface Network* (*PInet*) model. Predicted interaction probabilities are mapped onto the surface of G-actin and reported on a normalized 0–1 scale, with higher values indicating greater likelihood of nanocluster interaction.

**Supplementary References**


1. Shao, X. *et al.* Voltage Modulated Untwist Deformations and Multispectral Optical Effects from Ion Intercalation into Chiral Ceramic Nanoparticles. *Adv. Matererials* 10.1002.adma (2023).
2. Neese, F. Software update: The ORCA program system—Version 5.0. *WIREs Comput. Mol. Sci.* **12**, e1606 (2022).
3. Barone, V. & Cossi, M. Quantum Calculation of Molecular Energies and Energy Gradients in Solution by a Conductor Solvent Model. *J. Phys. Chem. A* **102**, 1995–2001 (1998).
4. Guilherme, L. R., Massabni, A. C., Dametto, A. C., de Souza Corrêa, R. & de Araujo, A. S. Synthesis, Infrared Spectroscopy and Crystal Structure Determination of a New Decavanadate. *J. Chem. Crystallogr.* **40**, 897–901 (2010).
5. Ramos, S., Moura, J. J. G. & Aureliano, M. Actin as a potential target for decavanadate. *J. Inorg. Biochem.* **104**, 1234–1239 (2010).
6. Sciortino, G., Aureliano, M. & Garribba, E. Rationalizing the Decavanadate(V) and Oxidovanadium(IV) Binding to G-Actin and the Competition with Decaniobate(V) and ATP. *Inorg. Chem.* **60**, 334–344 (2021).
7. Varadi, M. *et al.* AlphaFold Protein Structure Database in 2024: providing structure coverage for over 214 million protein sequences. *Nucleic Acids Res* **52**, D368–D375 (2024).
8. Jumper, J. *et al.* Highly accurate protein structure prediction with AlphaFold. *Nature* **596**, 583–589 (2021).
9. Wang, H., Robinson, R. C. & Burtnick, L. D. The structure of native G-actin. *Cytoskeleton* **67**, 456–465 (2010).
10. Dai, B. & Bailey-Kellogg, C. Protein interaction interface region prediction by geometric

deep learning. *Bioinformatics* **37**, 2580–2588 (2021).

11. Qi, C. R., Su, H., Mo, K. & Guibas, L. J. PointNet: Deep Learning on Point Sets for 3D Classification and Segmentation. Preprint at https://doi.org/10.48550/arXiv.1612.00593 (2017).
12. Cha, M. *et al.* Unifying structural descriptors for biological and bioinspired nanoscale complexes. *Nat. Comput. Sci.* **2**, 243–252 (2022).
13. Pierce, B. G., Hourai, Y. & Weng, Z. Accelerating protein docking in ZDOCK using an advanced 3D convolution library. *PLOS ONE* **6**, e24657 (2011).
14. Krishna, R. *et al.* Generalized biomolecular modeling and design with RoseTTAFold All-Atom. *Science* **384**, eadl2528 (2024).
15. Aureliano, M. & Crans, D. C. Decavanadate (V10O286-) and oxovanadates: Oxometalates with many biological activities. *J. Inorg. Biochem.* **103**, 536–546 (2009).
16. Gurel, P. S. *et al.* Cryo-EM structures reveal specialization at the myosin VI-actin interface and a mechanism of force sensitivity. *eLife* **6**, e31125 (2017).
